# AMPAR trafficking dependent LTP initiates cortical remapping and adaptive behaviors during sensory experience

**DOI:** 10.1101/2020.03.19.999094

**Authors:** Tiago Campelo, Elisabete Augusto, Nicolas Chenouard, Aron de Miranda, Vladimir Kouskoff, Daniel Choquet, Frédéric Gambino

## Abstract

Cortical plasticity improves behaviors and helps recover lost functions after injury by adapting neuronal computations. However, the underlying synaptic and circuit mechanisms remain unclear. In mice, we found that trimming all but one whisker enhances sensory responses from the spared whisker in the somatosensory barrel cortex and occludes whisker-mediated long-term potentiation (w-LTP) *in vivo*. In addition, whisking-dependent behaviors that are initially impaired by single whisker experience (SWE) rapidly recover when associated cortical regions remap. Blocking the surface diffusion of AMPA receptors (AMPARs) suppresses the expression of w-LTP in naïve mice with all whiskers intact, demonstrating that physiologically induced LTP *in vivo* requires AMPARs trafficking. We used this approach to demonstrate that w-LTP is required for SWE-mediated strengthening of synaptic inputs and initiates the recovery of previously learned skills during the early phases of SWE. Taken together, our data reveal that w-LTP mediates cortical remapping and behavioral improvement upon partial sensory deafferentation and demonstrates that restoration of sensory function after peripheral injury can be manipulated.

---

Functional sensory maps in the cerebral cortex reorganize in response to brain trauma or peripheral injury, with active modalities gaining cortical space at the expense of less active ones (Merzenich et al., 1983; Xerri, 2012). Map expansion has been proposed to adapt behaviors by optimizing neuronal circuits. Most of the evidence that map expansion is required for skills’ adaptation comes from studies that correlated behavioral changes with use-dependent map reorganization (Bieszczad and Weinberger, 2010; Molina-Luna et al., 2008; Reed et al., 2011). Despite these worthwhile contributions, the underlying circuit and synaptic mechanisms remain poorly understood. Here, we exploited the mouse whisker-to-barrel cortex system to explore the relationship between the synaptic mechanisms of sensory map plasticity and correlated adaptive behaviors.

Rodents use their whiskers to explore their immediate tactile environment. Under normal conditions, neurons in each barrel-column have receptive fields in the primary somatosensory cortex (S1) that are strongly tuned towards one principal whisker (PW). Nevertheless, trimming all but one whisker (SWE, single whisker experience) causes layer (L) 2/3 pyramidal neurons located in the deprived and spared-related columns to respond stronger to the spared whiskers stimulation (Feldman, 2009; Fox, 2002; Glazewski and Fox, 1996; Glazewski et al., 1996; Margolis et al., 2014), thereby resulting in the strengthening and expansion of the spared whisker representations within the map (Feldman, 2009; Fox, 2002; Margolis et al., 2014). Long-term synaptic potentiation (LTP) has been postulated as a synaptic mechanism for such response strengthening during learning and deprivation-induced plasticity (Bear et al., 1987; Clem et al., 2008; Feldman, 2009; Finnerty et al., 1999; Fox, 2002; Glazewski and Fox, 1996; Glazewski et al., 1996; Margolis et al., 2014; Rioult-Pedotti et al., 2000; Whitlock et al., 2006). In support of this idea, initial studies reported that activation of *N*-methyl-D-aspartate receptors (NMDARs) (Clem et al., 2008; Rema et al., 1998), α-amino-3-hydroxy-5-methyl-4-isoxazolepropionic acid receptors (AMPARs) (Clem and Barth, 2006; Dachtler et al., 2011), α/δ CREB, α-CaMKII, and α-CaMKII autophosphorylation (Glazewski et al., 1996, 2000) are all involved in neuronal response potentiation in L2/3. SWE increases synaptic strength of vertical and horizontal connections within and across cortical columns, and occludes electrically-induced, NMDAR-dependent LTP in acute brain slices obtained from young rodents (Clem and Barth, 2006; Clem et al., 2008; Finnerty et al., 1999), thereby providing consistent, yet indirect, evidence for a requirement of LTP during whisker map plasticity. However, while several paradigms to induce sensory-mediated LTP have been characterized *in vivo* (Gambino and Holtmaat, 2012; Gambino et al., 2014; Zhang et al., 2015), a direct demonstration that synaptic plasticity is required for cortical remapping and the adaptation of sensorimotor skills is still lacking. This likely results from 1) the technical difficulty to induce LTP in living animals with behaviorally-relevant stimuli; 2) the relatively limited knowledge on the molecular mechanisms underlying this form of plasticity *in vivo*, and 3) the lack of specific tools to block this LTP and spare the basal circuit functions while cortical regions remap.

Recently, manipulating the mobility of AMPARs has provided a new and specific tool to block LTP during behavior, without altering basal synaptic transmission and circuit functions (Humeau and Choquet, 2019; Penn et al., 2017). Here, we show that cross-linking GluA2 subunit *in vivo* inhibits LTP induced by physiological and behaviorally relevant stimuli (w-LTP), presumably by blocking GluA2 surface diffusion and preventing the increase in synaptic AMPAR content. This demonstrates, for the first time, a direct link between AMPAR trafficking and physiologically induced LTP *in vivo*, and reaffirms decades of *in vitro* works. Using this strategy, we demonstrate that w-LTP is causally inducing the potentiation of whisker-evoked responses and initiates behavioral recovery in the early phase of cortical remapping following whisker trimming. Altogether, our results highlight that the roles of LTP go beyond classic memory encoding functions: it also supports the adaptation of the cortical computation of perception, in particular in critical condition of sensory deprivation.

## Results

### SWE reshapes cortical sensory map and increases whisker-evoked responses

Previous studies have suggested that spared whiskers gain cortical space through LTP-mediated changes in the efficacy of existing synapses (Barth et al., 2000; Clem and Barth, 2006; Clem et al., 2008; Feldman, 2009; Feldman and Brecht, 2005; Glazewski and Fox, 1996; Glazewski et al., 1996). To explore this question *in vivo*, we first investigated the impact of single whisker experience (SWE) on whisker-evoked neuronal and synaptic responses. We exposed mice to a brief period of SWE (2-4 days) by clipping all but one C2 whisker and quantified the spatial representation of the spared whisker in contralateral S1 during anesthesia using intrinsic optical signal (IOS) imaging (**Fig. 1A-C**). For each mouse, IOS was acquired before, during, and after a 1-s long train (8 Hz) of C2 whisker deflection, in full-whisker experience (FWE, n=6) and SWE (n=7) animals. The whisker deflection-evoked response area was computed by using a pixel-by-pixel paired *t*-test between the averaged baseline and deflection-evoked IOS over at least 10 successive trials, as previously described (Schubert et al., 2013). In agreement with the potentiation of sensory-driven responses *in vivo* (Barth et al., 2000; Clem and Barth, 2006; Clem et al., 2008; Glazewski and Fox, 1996; Glazewski et al., 1996), the IOS evoked by the deflection of the spared whisker increased upon SWE within the spared whisker barrel column (**Fig. 1A-C**). Importantly, it occurred at a time at which no alterations in activity of layer 4 granular neurons have been observed in L2/3 (Glazewski and Fox, 1996; Glazewski et al., 1996), suggesting that SWE-induced map plasticity originates primarily from changes in neural activity within L2/3.

**Fig.1.**
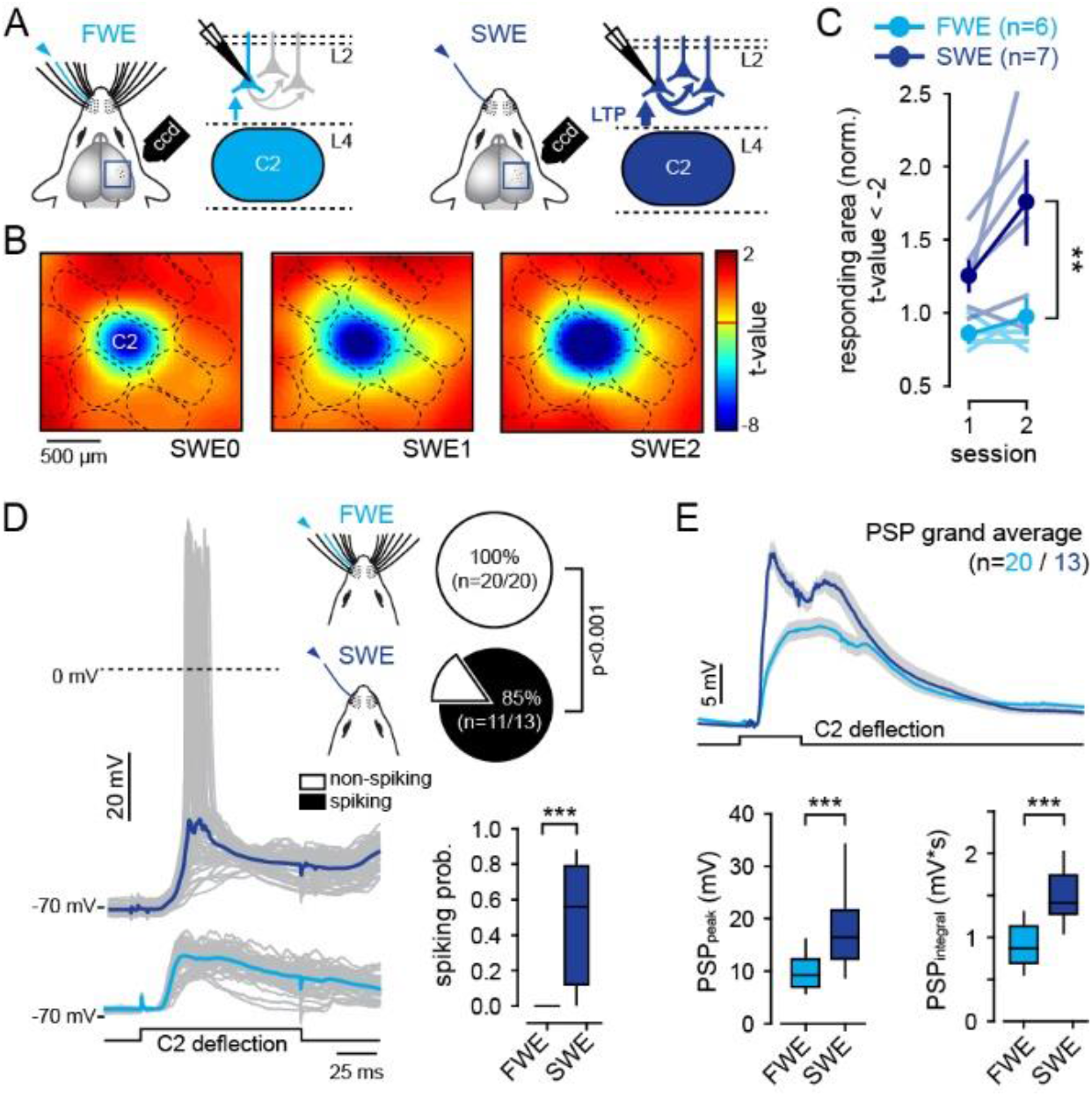
SWE increases whisker-evoked supra- and sub-threshold synaptic responses. **A)** Schematic of IOS and whole-cell recordings of L2/3 pyramidal neurons in full-whisker experience (FWE) and single-whisker experience (SWE) mice. PSPs IOS are evoked by deflecting the principal whisker (PW). **B)** Statistical *t*-maps over 3 successive days. For each mouse, red light reflectance 100 ms-long frames were acquired during anesthesia before (frames 1-10), during (frames 11-20), and after (frames 21-50) a 1-s long train (8 Hz) of single whisker deflection. The PW-evoked response area is computed by a statistical comparison of the averaged baseline (frames 1-10) and whisker-evoked (frames 19-28) IOS over at least 10 successive trials. This is done by using a pixel-by-pixel paired *t*-test, and only pixels with a *t*-value below the threshold (t<−2) are included into the stimulus-evoked response area. **C)** Averaged (± sem) PW-evoked response area (normalized to the first session at day 0). Light/dark blue lines, individual FWE/SWE mice, respectively. Circle, mean. **D)** *Left*, single-cell examples of whisker-evoked responses (grey, single trials; dark and light blue, averaged traces from SWE and FWE mice, respectively). Square pulse lines, C2 whisker deflection (100 ms). *Right*, fraction of spiking neurons (top) and number of spikes per whisker deflection (spiking probability; median ± interquartile range). **E)** *Top*, PW-evoked PSP grand average (all recorded cells averaged) ± sem. Square pulse lines, C2 whisker deflection (100 ms). *Bottom*, median (± interquartile range) PSP peak amplitude and integral. ***Values and statistical tests are provided in the Table S1*.**

We next performed whole-cell recordings of L2/3 pyramidal neurons *in vivo* in the C2 barrel while deflecting the principal (PW, C2) whisker, before and after SWE (**Fig. 1A, D, E**). In agreement with the above results, the fraction of spiking neurons in L2/3 and the number of spikes per PW deflection were increased in SWE as compared to FWE mice (fraction of spiking cells, FWE: 0/20; SWE: 11/13; p<0.001; spiking probability, FWE: 0; SWE: 0.48 ± 0.09; p<0.001) (**Fig. 1D**). To understand the mechanisms that alter the output of L2/3 neurons during SWE, we next examined whisker-evoked neuronal responses that did not generate spikes. As compared to FWE, SWE significantly increased the mean PW-evoked postsynaptic potential (PSP) peak amplitudes (FWE: 9.88 ± 0.86 mV; SWE: 17.98 ± 2.26 mV; p<0.001) (**Fig. 1E**). The concurrent increase in spiking probability and strengthening of synaptic transmission indicate that, despite a moderate increase in intrinsic excitability, the change of L2/3 neuronal spiking after SWE mostly resulted from an increase in peak amplitude of whisker-evoked subthreshold PSPs (**Fig. S1**).

### SWE occludes w-LTP

Because LTP-like mechanisms are prime candidates for enhancing synaptic transmission after SWE (Carvalho and Buonomano, 2009; Clem et al., 2008; Rioult-Pedotti et al., 2000; Whitlock et al., 2006), we next compared *in vivo* LTP induced by whisker stimulation (w-LTP) in FWE and SWE mice (**Fig. 2**). In FWE animals, a significant potentiation of whisker-evoked PSP was elicited by stimulating the PW for 1 min at a frequency of 8 Hz (RWS, rhythmic whisker stimulation) (baseline: 8.18 ± 1.17 mV, RWS: 9.77 ± 1.11 mV; n=7; p=0.002) (**Fig. 2B, C**, light blue traces; see also **Fig. 3D**). This potentiation was in good agreement with the w-LTP induced by RWS through NMDARs-dependent plateau potentials driven by the coordinated activation of segregated thalamo-cortical circuits (**Fig. 2A**) (Gambino et al., 2014; Zhang et al., 2015). In stark contrast, in SWE mice, RWS failed to strengthen whisker-evoked PSP (baseline: 20.45 ± 2.26 mV, RWS: 19.9 ± 2.12 mV; n=7; p=0.264) (**Fig. 2B, C**, dark blue traces). To investigate the possibility that the absence of w-LTP after SWE was the consequence of the alteration of its induction mechanism, we extracted and compared NMDARs-dependent plateau potentials in both FWE and SWE conditions, as previously described (Gambino et al., 2014). Compared to control FWE condition, plateau potentials evoked by single whisker stimulation were not significantly changed in SWE (plateau strength, FWE: 0.99 ± 0.03 mV*sec, n=20; SWE: 1.35 ± 0.08, n=13; p=0.3) (**Fig. 2E**). Neurons bearing high plateau potentials could not be potentiated, indicating that SWE has altered the positive correlation between plateau strength and the level of w-LTP observed in FWE mice (Gambino et al., 2014) (**Fig. 2F**, see also **Fig. S4**). This demonstrates that a key component of the NMDARs-dependent induction mechanism of w-LTP was not suppressed during SWE and could thus not account for the lower success rate of w-LTP. Instead, our results indicate that SWE enhances synaptic response to the spared whisker and occludes w-LTP (FWE: 123.5 ± 5.9 %, n=7; SWE: 97.6 ± 2.1 %, n=7; p=0.001) (**Fig. 2D-F**).

**Fig.2.**
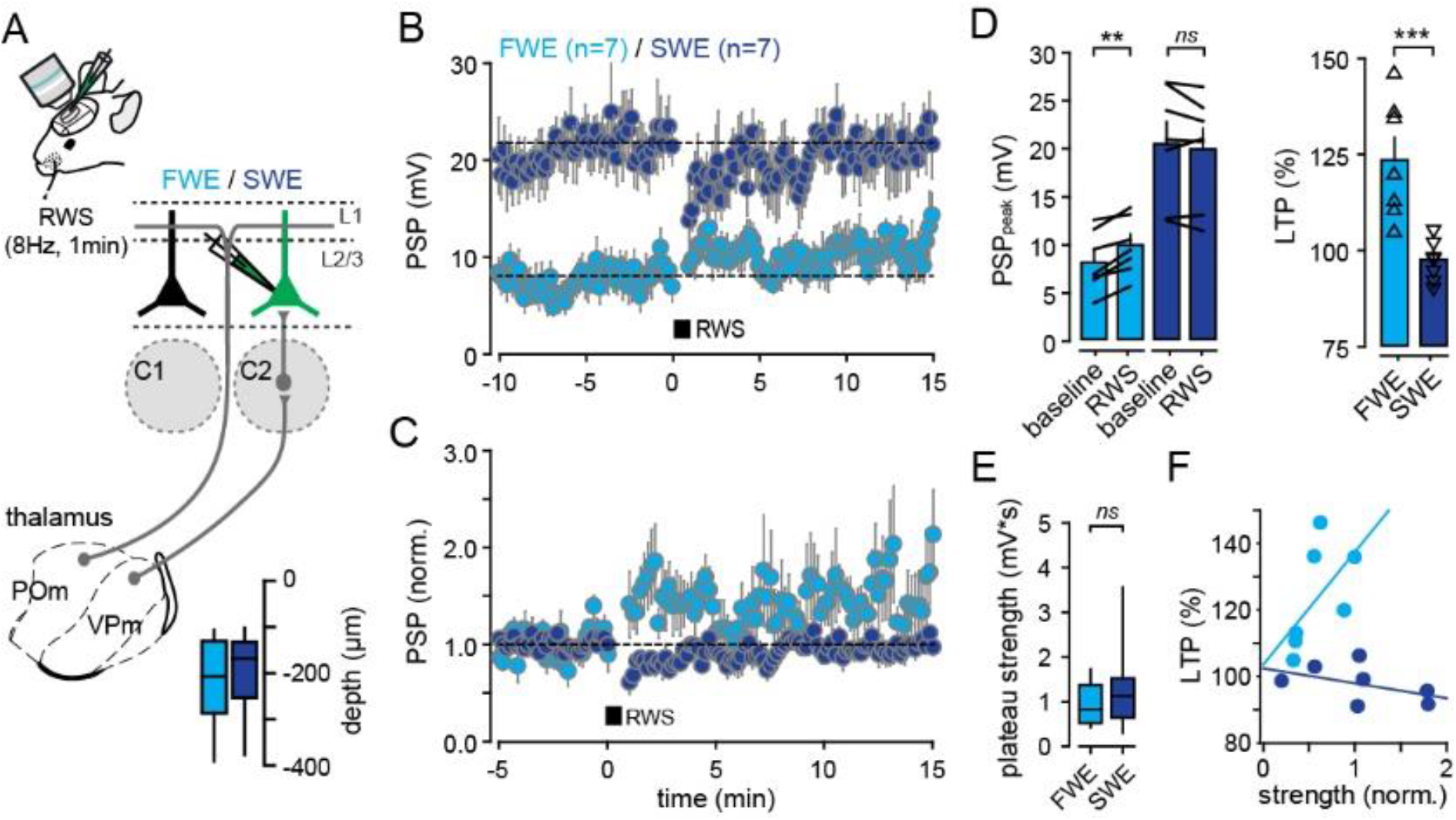
SWE occludes w-LTP. **A)** Schematic of thalamo-cortical circuits. Sensory information from the whiskers is transmitted to S1 by two main and well-segregated thalamo-cortical projections. L2/3 pyramidal neurons are recorded in the principal barrel-related column upon deflection of the PW (C2). Depth of recorded cells is indicated. **B)** Time-course of averaged PSP peak amplitude before and after RWS, in FWE and SWE mice. **C)** Time-course of averaged PSP amplitude normalized to baseline. **D)** *Left*, mean (±sem) amplitude before (baseline) and after RWS. Error bars, sem; black lines between bars, pairs. *Right*, mean (±sem) amplitude normalized to baseline (% of LTP). Triangles, individual cells. **E)** Median (± interquartile range) of plateau strength. **F)** Correlation between normalized plateau strength and the level of RWS-induced LTP in FWE (light blue) and SWE (dark blue) mice. SWE dissociates the induction from the expression of w-LTP by suppressing w-LTP without affecting plateau strength. ***Values and statistical tests are provided in the Table S1*.**

**Fig.3.**
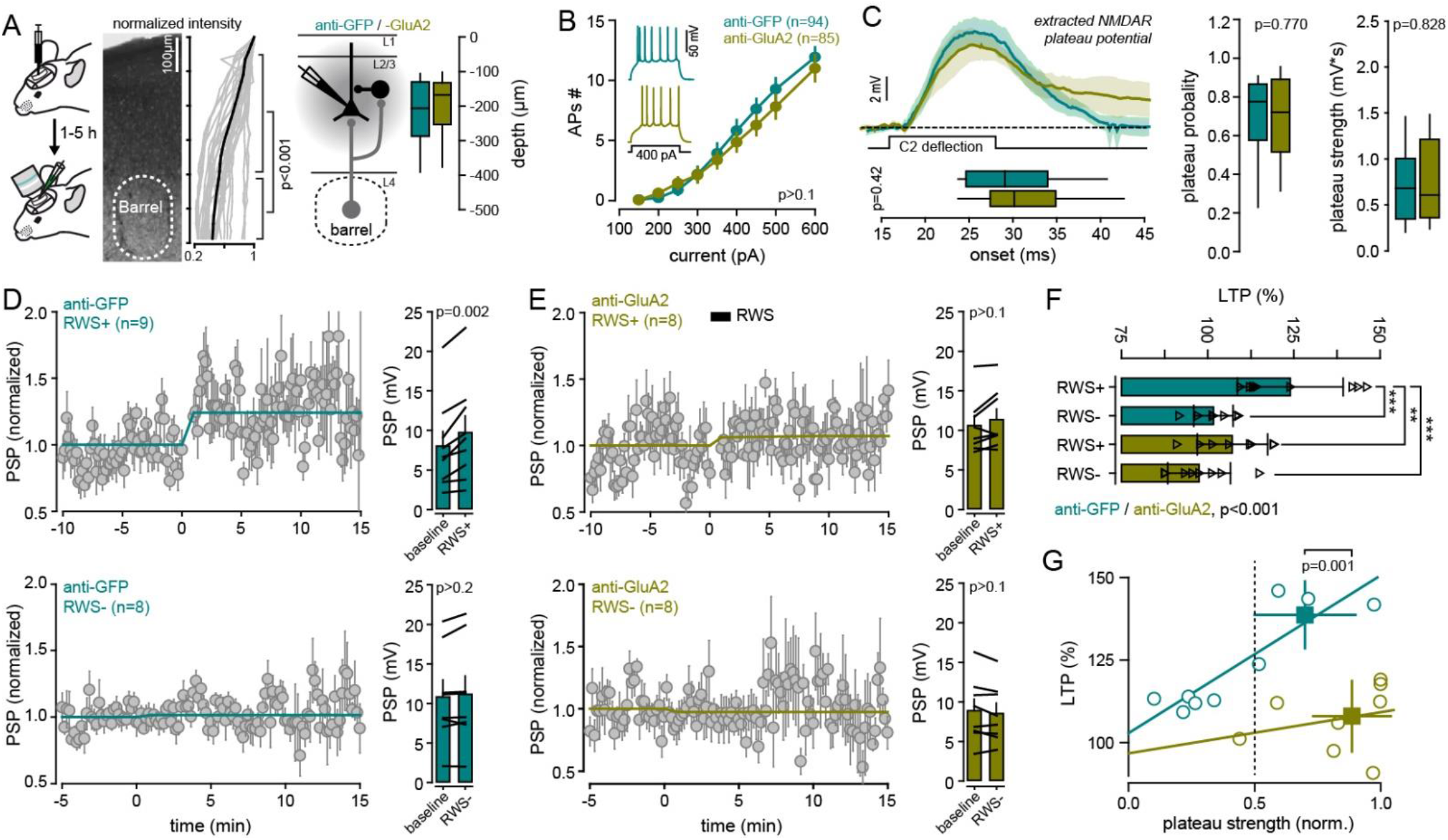
Cross-linking GluA2 subunit suppresses the expression of w-LTP without altering its induction mechanism. **A)** *Left*, schematic of experimental strategy. L2/3 pyramidal neurons are recorded in PW barrel-related column, 1 to 5h after IgGs injection. *Middle*, normalized intensity of anti-GluA2 IgGs signal as a function of cortical depth. IgGs are mostly targeting superficial layers. *Right*, schematic of the excitatory/inhibitory feed-forward circuit in a barrel-related column. Depth of recorded cells is indicated. **B)** Average (± sem) number of action potentials (APs) triggered by incremental current injections. *Insert*, example of spiking pattern in anti-GFP (green) and anti-GluA2 (yellow) injected mice upon 400pA current injection. **C)** *Left top*, grand average of PW-evoked extracted plateau potential (all recorded cells averaged ± sem). Black square pulse line, C2 whisker deflection (100 ms). *Left bottom*, median (± interquartile range) onset of plateau potentials. *Right*, median (± interquartile range) plateau probability and strength. **D)** *Left*, time-course of averaged PSP peak amplitude upon RWS (RWS+, *top*) and when RWS is not induced (RWS-, *bottom*), in anti-GFP injected mice. *Right*, mean (± sem) peak amplitude before (baseline) and after RWS+ (*top*) or RWS- (*bottom*). Black lines between bars, pairs. **E)** Same as in D) but for anti-GluA2 IgGs injected mice. **F)** Mean (± sem) peak amplitude normalized to baseline (% of LTP). Triangles, individual cells. **G)** Correlation between normalized plateau strength and the level of RWS-induced LTP in anti-GFP (green) and anti-GluA2 (yellow) IgGs injected mice. ***Values and statistical tests are provided in the Table S1.***

### Cross-linking AMPARs blocks the expression of w-LTP in FWE mice

Blocking the induction of LTP with NMDARs antagonists provided the most direct evidence that synaptic plasticity at appropriate synapses is required for both potentiation of spared whisker responses and learning (Clem et al., 2008; Rema et al., 1998; Takeuchi et al., 2013). However, NMDARs antagonists might obstruct normal sensory cortical transmission *in vivo*, which also relies on NMDARs conductances (Armstrong-James et al., 1993). Instead, we used an antibody cross-linking approach to limit the surface diffusion of postsynaptic GluA2 (Penn et al., 2017) and thus block the expression of LTP while sparing NMDARs conductances (Humeau and Choquet, 2019; Penn et al., 2017).

To investigate the role of AMPARs trafficking in LTP *in vivo* induced by whisker stimulation (w-LTP), we injected the immunoglobulins G (IgGs) against GluA2 (or the control anti-GFP IgG) in layer 2/3 of the right C2 cortical column, and performed whole-cell recordings in control FWE mice (**Fig. 3A**). We targeted this subunit of AMPAR as it is predominantly expressed in the neocortex (Schwenk et al., 2014), its expression in S1 is dynamically regulated upon partial sensory deafferentation (Gierdalski et al., 1999; He et al., 2004), and its immobilization has been shown to block LTP in acute slices and *in vivo* (Penn et al., 2017). First, to evaluate whether IgGs have an effect on basic cell and circuit electrophysiological properties, we recorded the firing properties, spontaneous membrane potential (Vm) fluctuation and whisker-evoked synaptic responses. No statistical differences in these parameters were detected between anti-GluA2 and anti-GFP IgGs in FWE mice (**Fig. 3B, C**; **Fig. S2** **and** **S3**). Consistent with previous studies (Humeau and Choquet, 2019; Penn et al., 2017), our results demonstrate that the cortical injection of anti-GluA2 IgGs does not affect intrinsic neuronal excitability, nor AMPARs- and NMDARs-mediated sensory-evoked synaptic transmission in basal conditions (**Fig. S2** **and** **S3**).

RWS induced a significant w-LTP of PW-evoked PSP in the presence of anti-GFP IgGs, similar to that observed in control non injected FWE mice (baseline: 8 ± 1.9 mV, RWS: 9.7 ± 2 mV; n=9; p=0.002) (**Fig. 3D**). On average, the change in PSP amplitude when RWS was applied (RWS+: 123.9 ± 1.7 %, n=9) was significantly higher than when RWS was not (RWS−: 101.6 ± 0.71 %, n=8, p<0.001) (**Fig. 3F**), and positively correlated with the strength of plateau potentials (**Fig. 3G**; **Fig. S4**). In contrast, cross-linking-mediated suppression of GluA2 diffusion prevented w-LTP (baseline: 10.6 ± 1.2 mV, RWS: 11.3 ± 1.3 mV; n=8; p=0.102; RWS+ *vs.* RWS−: 107.1 ± 3.6 % *vs.* 97.5 ± 3.1 %, p>0.05) (**Fig. 3E, F**; **Fig. S4**). Compared to control IgGs, anti-GluA2 IgGs did not significantly change NMDARs-dependent plateau potentials evoked by single whisker stimulation (plateau probability: anti-GFP: 0.70 ± 0.04, n=26; anti-GluA2: 0.68 ± 0.05, n=24; p=0.770; plateau strength: anti-GFP: 0.71 ± 0.09 mV*sec, n=26; anti-GluA2: 0.77 ± 0.11 mV*sec, n=24; p=0.828) (**Fig. 3C**). S1 pyramidal neurons bearing high plateau strength could not be potentiated in the presence of anti-GluA2 IgGs (plateau strength > 0.5; anti-GFP: 138.6 ± 5 %, n=4; anti-GluA2: 108 ± 4 %, n=7; p<0.001) (**Fig. 3G**; **Fig. S4**). Altogether, these results demonstrate that cross-linking surface GluA2 *in vivo* prevents the expression of w-LTP, but not the NMDARs-dependent plateau potentials responsible for induction of w-LTP in control animals, thereby complementing *in vitro* observations (Choquet, 2018; Granger et al., 2012; Penn et al., 2017).

### w-LTP mediates neuronal potentiation during SWE-induced cortical remapping

Our data demonstrate that, as opposed to the pharmacological blockade of NMDARs, cross-linking GluA2 subunits represents an effective and straightforward way to prevent the expression of w-LTP *in vivo* without modifying its induction mechanisms or basal synaptic transmission. Thus, we used this approach to question if w-LTP was causally inducing the potentiation of whisker-evoked response during SWE. We reasoned that if w-LTP increases synaptic responses during SWE, the chronic suppression of GluA2 surface diffusion during SWE would block this mechanism, thereby allowing RWS to potentiate whisker-evoked PSP when GluA2 mobility is restored.

To test these predictions, anti-GluA2 IgGs (or anti-GFP for controls) were injected in S1 twice a day for two consecutive days while trimming all but the contra-lateral C2 whisker. L2/3 pyramidal neurons were then recorded after a 12h-clearance period to washout IgGs (X-SWE, **Fig. 4A**). X-SWE significantly decreased the average number of spikes per PW deflection (anti-GluA2: 0.08 ± 0.03; n=9; anti-GFP: 0.35 ± 0.14, n=8; p=0.026) (**Fig. 4B, C**) although it did not modify the fraction of spiking neurons (X-SWE: 6/9; SWE: 11/13; p>0.05) (**Fig. 4B**). The average PW-evoked PSP peak amplitude in the presence of anti-GluA2 IgGs (7 ± 1.3 mV, n=9, p<0.001), but not anti-GFP IgGs (14.2 ± 1.6 mV, n=8, p=0.126), was significantly decreased as compared to SWE (17.9 ± 2.3 mV, n=13) (**Fig. 4D, E**). X-SWE did not alter whisker-induced plateau potentials (**Fig. S4B, C**). RWS potentiated PW-evoked PSP when anti-GluA2 IgGs, but not anti-GFP IgGs, were washed-out (anti-GluA2; baseline: 7.6± 1.7 mV, RWS: 10.4± 2.6 mV; n=6; p=0.04; anti-GFP; baseline: 12.6 ± 1.6 mV, RWS: 12.2 ± 1.9 mV; n=6; p=0.436) (**Fig. 4F, G**). Thus X-SWE with anti-GluA2 IgGs preserved the expression of w-LTP in SWE mice (X-SWE: 129.8 ± 7.8 %, n=6 *vs*. SWE: 97.6 ± 2.1 %, n=7; p=0.001) to similar levels as in FWE mice (123.5± 5.9 %, n=7; p=0.4) (**Fig. 4G**; **Fig. S4**). This indicates that chronically blocking AMPARs trafficking during SWE prevents sensory-evoked synaptic potentiation.

**Fig.4.**
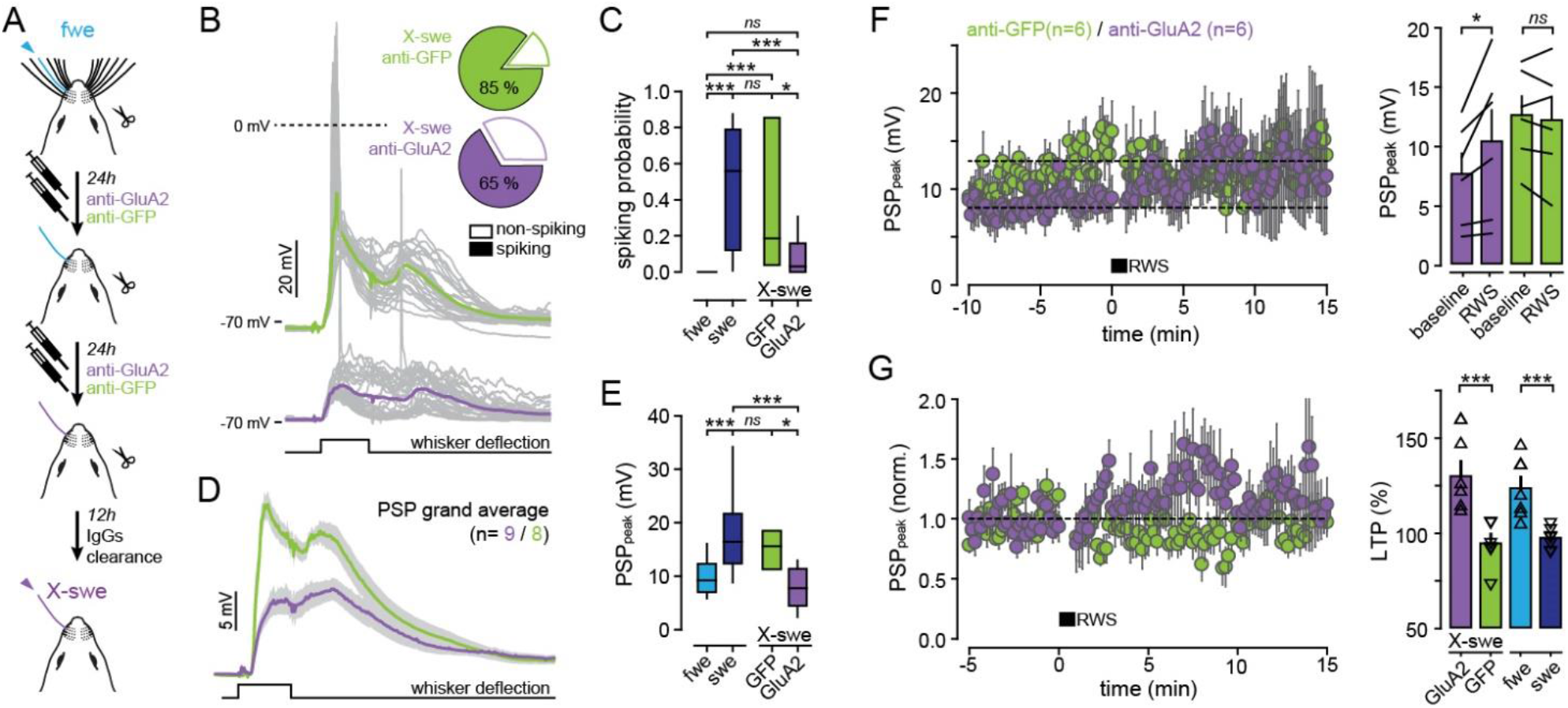
w-LTP mediates neuronal potentiation during SWE-induced cortical remapping. **A)** Schematic of experimental strategy. IgGs are injected during SWE, followed by washed-out before recordings. **B)** *Left*, single-cell examples of whisker-evoked responses (grey, single traces; green and purple, averaged traces from anti-GFP and anti-GluA2 injected mice, respectively). Square pulse line, C2 whisker deflection (100 ms). *Right*, fraction of spiking neurons triggered by whisker deflection. **C)** Number of spikes per whisker deflection (spiking probability; median ± interquartile range). **D)** PW-evoked PSP grand average (all recorded cells averaged ± sem). Square pulse line, C2 whisker deflection (100 ms). **E)** Median (± interquartile range) PSP peak amplitude. **F)** *Left*, time-course of averaged PSP peak amplitude upon RWS in anti-GFP (green) and anti-GluA2 (purple) injected mice (after wash-out). *Right*, mean (± sem) PSP peak amplitude before (baseline) and after RWS. Black lines between bars, pairs. **G)** *Left*, time-course of averaged PSP amplitude normalized to baseline. *Right*, mean (±sem) amplitude normalized to baseline (% of LTP). Triangles, individual cells. ***Values and statistical tests are provided in the Table S1.***

Importantly, there was no differences in basal synaptic transmission between SWE and X-SWE in presence of the control anti-GFP IgGs (**Fig. 4B-E**; **Fig. S4B**), suggesting that the chronic presence of antibodies did not alter receptors functions (Haselmann et al., 2018; Peng et al., 2014). Nevertheless, to exclude the possibility that chronic exposure to AMPARs cross-linking antibodies caused inflammation that could have affected cell and circuit properties, we performed immunohistochemical detection of astrocyte and microglia cell markers (**Fig. S5**). We injected anti-GluA2 and anti-GFP IgGs in the right and left hemispheres respectively, twice a day for two consecutive days, and assessed the number of astrocytes and microglia using anti-GFAP and anti-Iba1 immunostaining (**Fig. S5**). We found no quantitative differences between groups as the number of immunopositive cells as well as the total intensity of staining in the left and right hemisphere were similar, indicating that the astrocytic and microglial responses following injection was not potentiated in the presence of anti-GluA2 IgGs. In addition, the global locomotor activity appeared unaffected in mice chronically injected with IgGs in S1 (see **Fig. S7**), revealing that the prolonged cross-linking of AMPARs has no cytotoxic effect. Taken together, our results indicate that sensory LTP is thus causally inducing the potentiation of whisker-evoked response following whisker trimming.

### w-LTP facilitates the recovery of altered whisker-dependent behaviors during the early phases of SWE

We demonstrated above that the chronic blockade of AMPAR trafficking prevented potentiation of whisker-evoked responses during SWE, supporting the idea that w-LTP contributes to SWE-induced cortical remapping. Because SWE alters various whisker-mediated behavioral tasks (Barnéoud et al., 1991; Celikel and Sakmann, 2007; Clem et al., 2008; Xerri, 2012), we reasoned that if cortical remapping improves tactile perception, blocking w-LTP during SWE should affect whisker-mediated behavioral performance.

To test this hypothesis, we monitored freely behaving mice performing a binary gap-crossing task under infrared light (**Fig. 5A**; **Fig. S6A-C**). Mice were trained to reach a rewarding platform separated by a randomized distance between 40 and 65 mm from the home platform (**Fig. 5B**). At a distance of 65 mm, mice used preferentially their whiskers to locate the target platform and jump onto it to receive the reward (Barnéoud et al., 1991; Celikel and Sakmann, 2007) (**Fig. S6D**). SWE was induced after mice reached expertise (4 days of training) (**Fig. 5B**; **Fig. S6C**). Gap-crossing performance decreased immediately after SWE (fraction of success; session 4: 0.96 ± 0.04; session 5: 0.68 ± 13; n=6; p=0.006) but recovered quickly after 2 days of SWE (session 6: 0.87 ± 0.09; p=0.372) (**Fig. 5C**), a time scale at which whisker-evoked responses saturated (**Fig. 1**) and w-LTP has been fully occluded (**Fig. 2**). Importantly, mice that were not tested during SWE (sessions 5 to 7) had similar final success rate (session 8; 0.95 ±0.03, n=6; 0.89 ± 0.08, n=5; p>0.05) (**Fig. 5C, D**), suggesting that behavioral recovery was likely not caused by a new learning phase but instead resulted from the restoration of accurate perception. The gap-crossing performance of anti-GluA2 IgGs-injected mice decreased more (session 5: −61.8 ± 12% *vs.* −22.6 ± 6.7%, p=0.016) and recovered significantly slower as compared to that of anti-GFP injected mice (**Fig. 5E-G**). Success rates were however similar between both groups 3 days after SWE (session7: 0.82 ± 0.09 *vs.* 0.72 ± 0.14; p=0.805) (**Fig. 5G**), which might reflect barrel cortex-independent behavioral strategies (Celikel and Sakmann, 2007; Hong et al., 2018) and/or the existence of mechanisms that preserve a slow capacity for cortical remapping (Clem et al., 2008). None of the IgGs altered exploration and decision latency (**Fig. S7**). Altogether, our data indicate that blocking GluA2 diffusion similarly affects neuronal response potentiation *in vivo* and behavioral output at early phases of SWE, thereby providing new evidence for a critical role of w-LTP in facilitating the recovery of lost perceptual skills.

**Fig.5.**
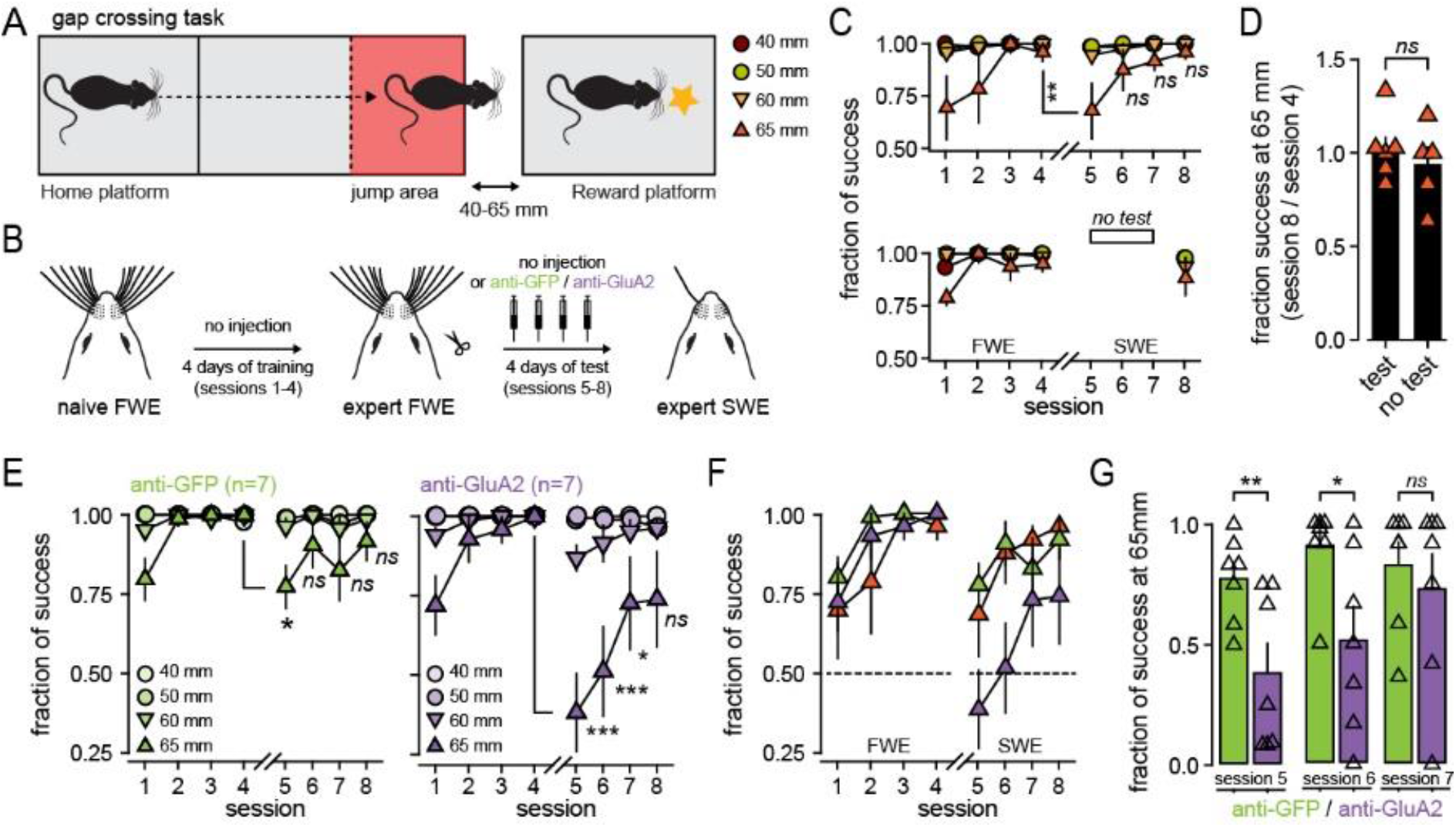
w-LTP facilitates the recovery of altered whisker-dependent behaviors during the early phases of SWE. **A)** Overview of the gap-crossing task (see extended data Fig. 4 for details). The reward platform is moved between trials to set the gap width from 40 to 65 mm. B) Schematic of the time-course regarding the behavior, the trimming of the whiskers and IgGs injections. Mice learn to reach the rewarding platform (4 consecutive days) before SWE is induced during which anti-GFP or anti-GluA2 IgGs are injected through implanted cannula twice a day (before and after each behavioral session). **C)** *Top*, averaged (±sem) fraction of gap-crossing success for different gap distances, in non-injected mice. *Bottom*, tests in session 5 to 7 were omitted to assess the role of learning during SWE. **D)** Mean (± sem) fraction of success in the final session (normalized to session 4 before SWE) at a distance of 65 mm for mice that are tested every day (test) and for mice that were not tested in sessions 5 to 7 (no test). Triangles, individual mice. **E)** Averaged (± sem) fraction of gap-crossing success for different gap distances, in anti-GFP (*left*) and anti-GLuA2 (*right*) injected mice. **F)** Averaged (± sem) fraction of gap-crossing success at a distance of 65 mm, in non-injected (orange), anti-GFP (green) and anti-GLuA2 (purple) injected mice. **G)** Mean (± sem) fraction of success at a distance of 65 mm after expertise in FWE mice and during SWE. Triangles, individual mice. ***Values and statistical tests are provided in the Table S1.***

## Discussion

Our study provides the best evidence to date that the rules governing LTP *in vitro* through the synaptic regulation of GluA2/GluA1 AMPARs (Granger et al., 2012; Penn et al., 2017; Shi et al., 2001) are effectively used *in vivo* and mediate the potentiation of spared inputs during cortical remapping. While the surface diffusion and insertion of AMPARs into the postsynaptic membrane has become a well-recognized hallmark of NMDAR-dependent synaptic potentiation *in vitro* (Diering and Huganir, 2018; Humeau and Choquet, 2019), the molecular mechanisms of synaptic LTP *in vivo* remained poorly unexplored. Here, we found that whisker-mediated LTP (w-LTP) could not readily be produced in S1 pyramidal neurons when rhythmic whisker stimulation (RWS) was applied in presence of anti-GluA2 IgGs (**Fig. 3D-F**). This is in agreement with earlier study showing that RWS triggers the insertion of AMPARs at dendritic spines in the barrel cortex in vivo (Zhang et al., 2015), and confirms that GluA2/A1 heteromers are the dominant form of AMPARs in the barrel cortex of adult animals (Kondo et al., 1997; Schwenk et al., 2014). Together, our data indicate that blocking the traffic of endogenous GluA2-containing AMPARs is sufficient to inhibit the expression of this form of LTP while leaving its induction mechanism (**Fig. 3C, G**) as well as the basal synaptic transmission unaltered (**Fig. S2** **and** **S3**).

We further exploited this result to describe the relationship between synaptic plasticity and cortical remapping following single-whisker experience (SWE). In acute brain slices, SWE has been shown to occlude electrically-induced, NMDAR-dependent LTP (Clem et al., 2008), but whether LTP occurs *in vivo* remained unknown. Here, we found that sensory inputs potentiated during SWE could not be further potentiated by RWS, indicating that SWE occludes RWS-mediated LTP (**Fig. 2**). This is the first direct demonstration, to our knowledge, that LTP induced *in vivo* by behaviorally-relevant stimuli (Gambino et al., 2014) occurs in S1 during SWE, which might drive the spared whisker representation in S1 to strengthen and broaden into deprived columns (**Fig 1B**). Consistent with this idea, SWE potentiates neuronal response to spared whisker deflections *in vivo* (**Fig. 1D-E**) (Barth et al., 2000; Glazewski et al., 1996, 2000; Rema et al., 1998), and enhances synaptic communication between L4-to-L2/3 and L2/3-to-L2/3 neurons *in vitro* (Cheetham et al., 2007; Clem and Barth, 2006; Clem et al., 2008; Finnerty et al., 1999). However, it remained unclear whether LTP is causally inducing the potentiation of whisker-evoked response following whisker trimming.

To address this question, we chronically cross-linked AMPAR during SWE (X-SWE), and tested the causal relation between w-LTP and SWE-induced cortical remapping. We found that X-SWE reduces the AMPAR-component of whisker-evoked PSP to levels similar as in naïve FWE mice (**Fig. 4B-E**), and restores the capacity to undergo w-LTP (**Fig. 4F, G**) after IgGs wash-out. It indicates that inhibiting the synaptic delivery of endogenous AMPARs prevents the potentiation of spared whisker-evoked PSP that would have otherwise occurred during SWE. Our results demonstrate, for the first time, that w-LTP, through the tight modulation of GluA2-containing AMPARs, is one of the major mechanisms that potentiates neuronal response during cortical remapping. We confirmed that our results were not caused by the alteration of receptor function, distribution and basal transmission, which might occur in the prolonged presence of certain types of anti-AMPARs antibodies (Haselmann et al., 2018; Peng et al., 2014). In addition to our controls of basal synaptic transmission (**Fig. S2 and S3**), we measured and compared the astrocytic/microglia response, NMDARs-dependent plateau potentials, and the global motor activity in presence of anti-GluA2 and anti-GFP IgGs. None of these parameters were different between conditions, indicating that basic cellular, circuit and behavioral properties were preserved during X-SWE (**Fig. S5** **and** **S7**). Taken together, our results are in line with the dynamic regulation of GluA2 expression in S1 upon partial sensory deafferentation (Gierdalski et al., 1999; He et al., 2004). However, they stand in contrast to the traditional view of AMPAR trafficking during cortical remapping, which states that GluA1 rather than GluA2 subunit is required for experience-dependent plasticity in the barrel cortex (Feldman, 2009; Makino and Malinow, 2011). Consistent with this model, previous *in vitro* studies in the barrel cortex of ~P15 animals concluded that SWE drives LTP at L4-L2/3 synapses by inserting homomeric GluA1 (GluA2-lacking) AMPARs (Clem and Barth, 2006; Clem et al., 2008; Miyazaki et al., 2012; Takahashi et al., 2003). Nevertheless, the GluA1 dependence of cortical and hippocampal plasticity has been shown to be developmentally regulated and decreases with age (Grosshans et al., 2002; Jensen et al., 2003). In addition, neuronal response potentiation is only partially blocked in older GluA1 knockout animals undergoing SWE (Hardingham and Fox, 2006), supporting the idea that GluA2/GluA1-dependent LTP indeed operates during neuronal response potentiation in adult animals.

Although our results demonstrate a tight relation between RWS-LTP and cortical remapping, how such local synaptic changes contribute to overall behavioral adaptation upon partial sensory deafferentation remains unknown. A first attempt in addressing this issue comes from pharmacological experiments where NMDARs antagonists were used during SWE, assuming they were selective for plasticity and not normal information processing in the brain (Clem et al., 2008; Rema et al., 1998). However, NMDARs antagonists have strong attenuating effect on long-latency spikes in supragranular layers, as well as in granule cell layers (Armstrong-James, 1993), which might be driven by dendritic plateau potentials generated by non-linearly summating local and thalamo-cortical inputs (Gambino et al., 2014; Lavzin et al., 2012; Palmer et al., 2014). Instead, blocking AMPARs trafficking with cross-linking antibodies has great potential to address this question *in vivo* (Humeau and Choquet, 2019). The content of AMPARs at synapses *in vivo* has be shown to correlate with motor performance (Roth et al., 2019), and blocking AMPARs trafficking impairs learning (Penn et al., 2017). Here, by manipulating the mobility of GluA2, we found that chronic cross-linking slows down behavioral recovery after SWE (**Fig. 5E-G**) while it did not alter sensory processing (**Fig. S7**). Strikingly, success rates were not different between anti-GFP and anti-GluA2 injected mice 3 days after SWE. This suggests that w-LTP is associated with a fast adaptation of cortical processing and related behaviors to the change of perceptual condition, while other adaptation mechanisms, such as mGluR-mediated synaptic plasticity (Clem et al., 2008), disinhibition (Isaacson and Scanziani, 2011) or homeostatic scaling (Zhang and Linden, 2003), could also be at play. For example, whisker trimming increases the pruning of inhibitory synapses (Chen et al., 2011; Keck et al., 2011) and weakens feed-forward inhibition in S1 L2/3 pyramidal neurons (House et al., 2011; Jiao et al., 2006). Such disinhibition could facilitate synaptic LTP during SWE (Gambino and Holtmaat, 2012; Williams and Holtmaat, 2019), for example by promoting dendritic signaling (Gambino et al., 2014).

Taken together, our results demonstrate that w-LTP initiates cortical remapping and occurs nearly immediately following partial sensory deafferentation, thereby providing new important processing resources for spared inputs that compensate for the loss of surroundings inputs. In support of this hypothesis, training-related increases in cortical representations correlate with perceptual learning (Bieszczad and Weinberger, 2010; Molina-Luna et al., 2008; Reed et al., 2011), suggesting that sensory deafferentation could cause behavioral gains by promoting cortical remapping. Our results are of importance for clinicians and patients, as brief periods of sensory deprivation have been proposed as therapeutic ways to promote recovery of lost function after peripheral injury or stroke (Kraft et al., 2018).

## Acknowledgements

We thank E. Normand and the IINS *in vivo* core facility for animal husbandry. We thank the IINS cell biology core facilities (LABEX BRAIN [ANR-10-LABX-43]) and in particular C. Breillat for antibody handling. We thank H. El Oussini and B. Darracq (Imetronic) for their technical expertise and support, and all the members of the Gambino and Choquet laboratories for technical assistance and helpful discussions. We thank E. Gouaux for generous gift of the 15F1 anti-GluA2 antibody.

This project has received funding from (to FG): the European Research Council (ERC) under the European Union’s Horizon 2020 research and innovation program (NEUROGOAL, grant agreement n° 677878), the FP7 Marie-Curie Career Integration program (grant agreement n° 631044); the ANR JCJC (grant agreement n° 14-CE13-0012-01), the University of Bordeaux (Initiative of Excellence senior chair 2014); (to DC): the European Research Council (ERC) under the European Union’s Horizon 2020 research and innovation program (ADOS, grant agreement n° 339541; DynSynMem, grant agreement n° 787340); and from the Region Nouvelle Aquitaine to DC and FG.

## Author contribution

TC, EA, AM, VK performed the experiments. TC, DC and FG conceived the studies and analyzed the data with the help of EA, NC, AM and VK. DC and FG supervised the research and wrote the manuscript with the help from co-authors.

## Extended Data Figures and Legends

**Fig. S1.**
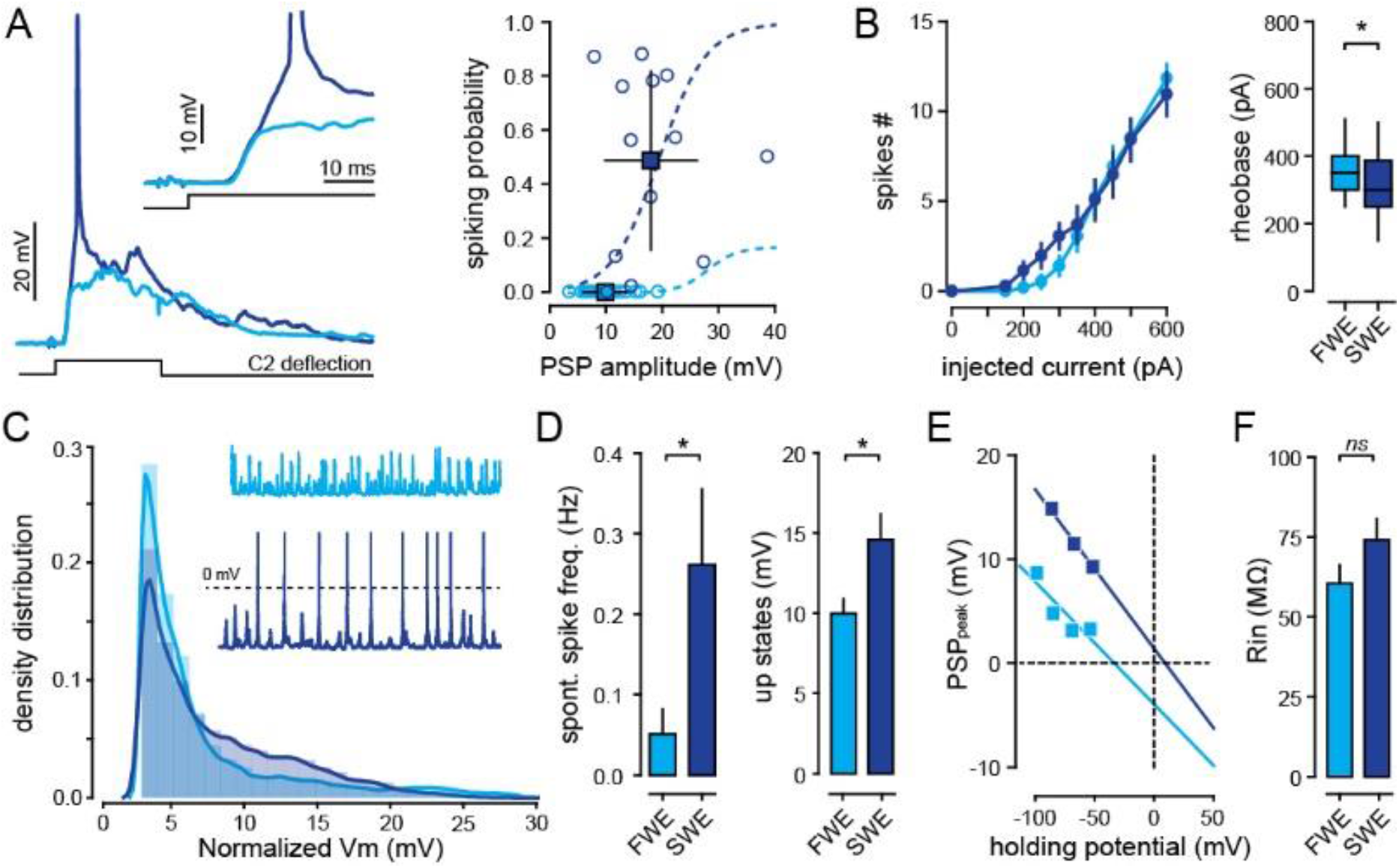
Effect of SWE on L2/3 pyramidal neurons excitability. **A)** *Left*, single-cell examples of PW-evoked responses (averaged traces from FWE (light blue) and SWE (dark blue) mice). *Right*, relationship between PW-evoked PSP amplitude and the spiking probability illustrating the increase in PSP-spike coupling upon SWE. Circles, individual cells; squares, average. **B)** *Left*, average (± sem) number of action potentials (APs) triggered by incremental current injections in FWE (light blue) and SWE (dark blue) mice. *Right*, Median (± interquartile range) minimal current amplitude (pA) triggering action potentials (rheobase). **C)** Averaged spontaneous membrane potential probability histograms. FWE, light blue; SWE, dark blue. *Insert*, single cell examples. **D)** *Right*, mean (± sem) frequency of spontaneous action potentials. *Left*, mean (± sem) amplitude of spontaneous up-states. FWE, light blue; SWE, dark blue. **E)** Correlation between holding membrane potential and whisker-evoked PSP amplitude in down-states. The reversal potential is higher in SWE (dark blue) as compared to FEW (light blue) suggesting disinhibition. **F)** Mean (± sem) membrane resistance. FWE, light blue; SWE, dark blue. ***Values and statistical tests are provided in the Table S1***.

**Fig. S2.**
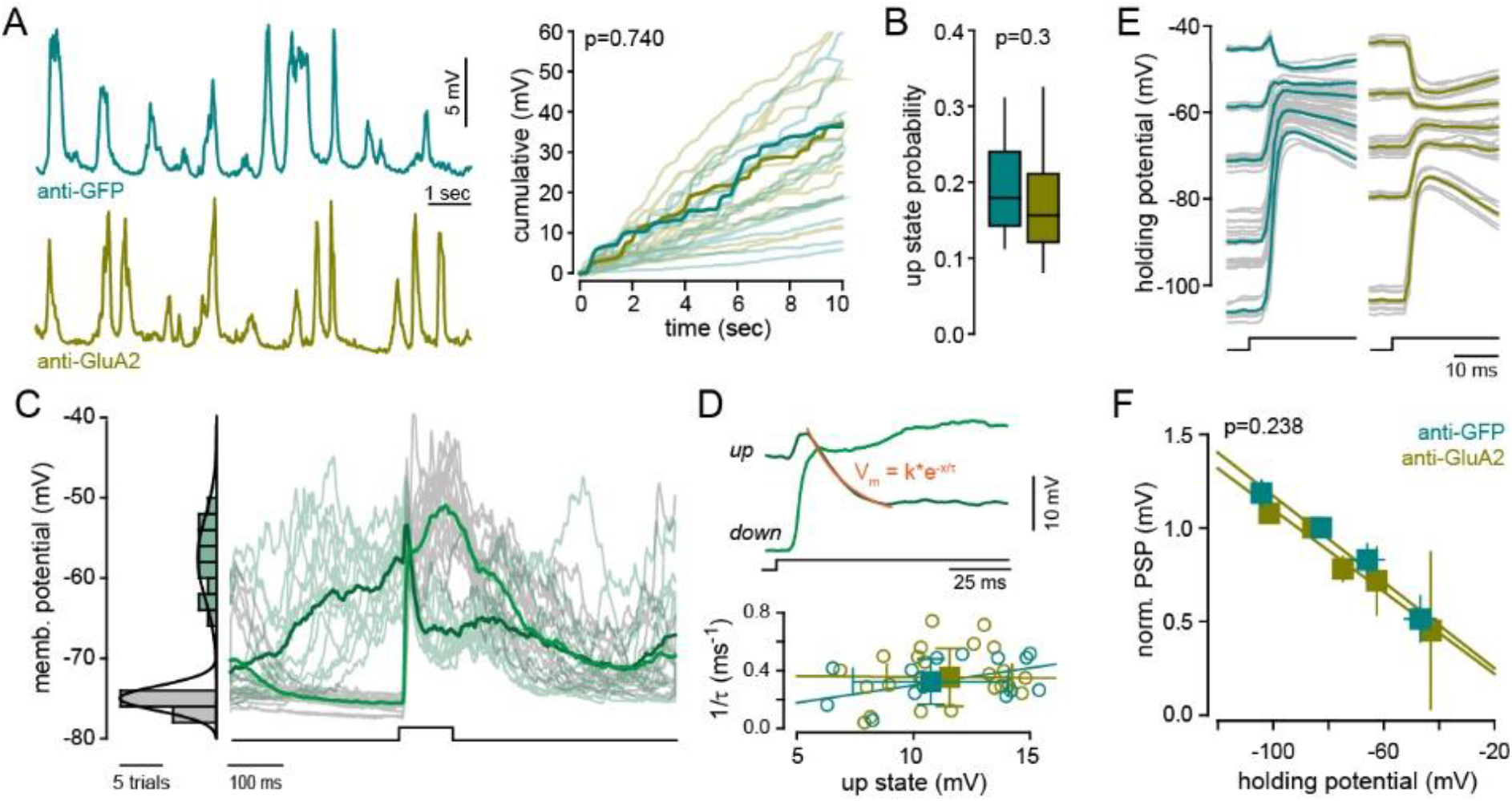
Cross-linking GluA2 subunit does not affect spontaneous activity nor PW-evoked inhibitory responses. **A)** *Left*, examples of single-cell spontaneous membrane potential during anesthesia in anti-GFP (top) and anti-GluA2 (bottom) IgGs injected mice. *Right*, cumulative RWS-induced depolarization. Light lines, individual cells. Bold lines, examples from left. **B)** Median (± interquartile range) probability of spontaneous up-states. **C)** *Left*, membrane potential histogram showing the average (30 ms) membrane potential before each PW stimulation. Down (grey) and up (green) states follow separated Gaussian distributions. *Right*, PW-evoked PSPs during down (green) and up (dark green) states. Individual trials are represented with light lines. **D)** *Top*, single-cell examples of PW-evoked PSP in down and up states. The decay of membrane potential during up states is fitted with an exponential, which is indicative of the amount of PW-evoked inhibition. Bottom, relation between the amplitude of up states and the exponential tau, in anti-GFP (green) and anti-GluA2 (yellow) IgGs injected mice. Circles, individual cells; squares, mean (± sem). **E)** single-cells examples of PW-evoked PSPs at different holding potentials. **F)** Relation between holding potential and the amplitude of PW-PSP (normalized to the amplitude at resting membrane potential). ***Values and statistical tests are provided in the Table S1.***

**Fig. S3.**
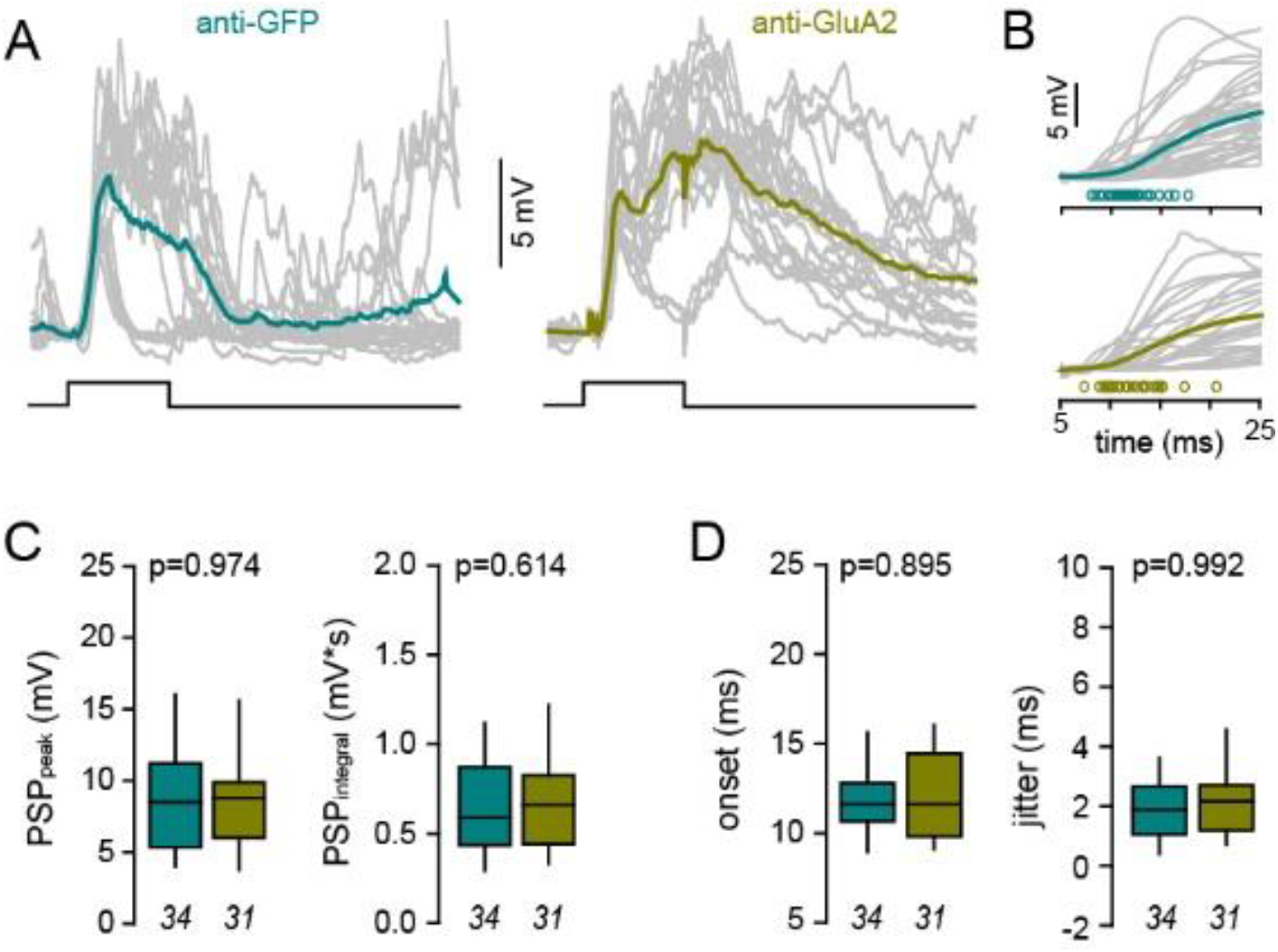
Cross-linking GluA2 subunit does not affect PW-evoked excitatory responses. **A)** *S*ingle-cell example of PW-evoked PSPs. Individual trials are represented with grey lines. Square pulse line, whisker deflections (100 ms). **B)** Single-cell examples of PW-evoked PSPs illustrating the onset of PSP. Circles, individual cells. **C)** Median (± interquartile range) PSP peak amplitude and integrals. **D)** Median (± interquartile range) PSP onset and onset jitter. ***Values and statistical tests are provided in the Table S1.***

**Fig. S4.**
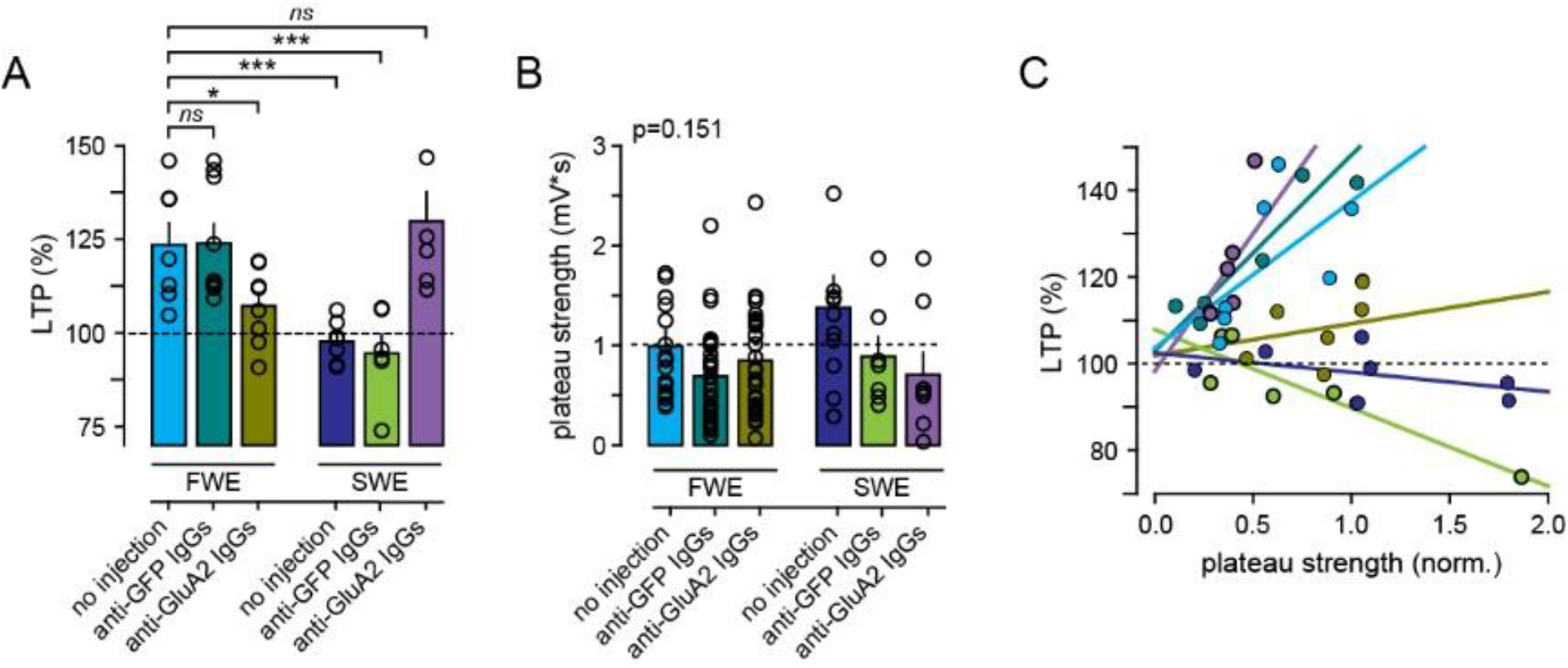
Comparison between all the different treatments for LTP and plateau strength. **A)** Mean (± sem) amplitude normalized to baseline (% of LTP). Circles, individual cells. **B)** Mean (± sem) plateau strength. Circles, individual cells. **C)** Correlation between normalized plateau strength and the level of RWS-induced LTP for all treatments. Only the conditions SWE (dark blue) and FWE+antiGluA2 IgGs (yellow) dissociate the induction from the expression of w-LTP by suppressing w-LTP without affecting plateau strength. ***Values and statistical tests are provided in the Table S1*.**

**Fig. S5.**
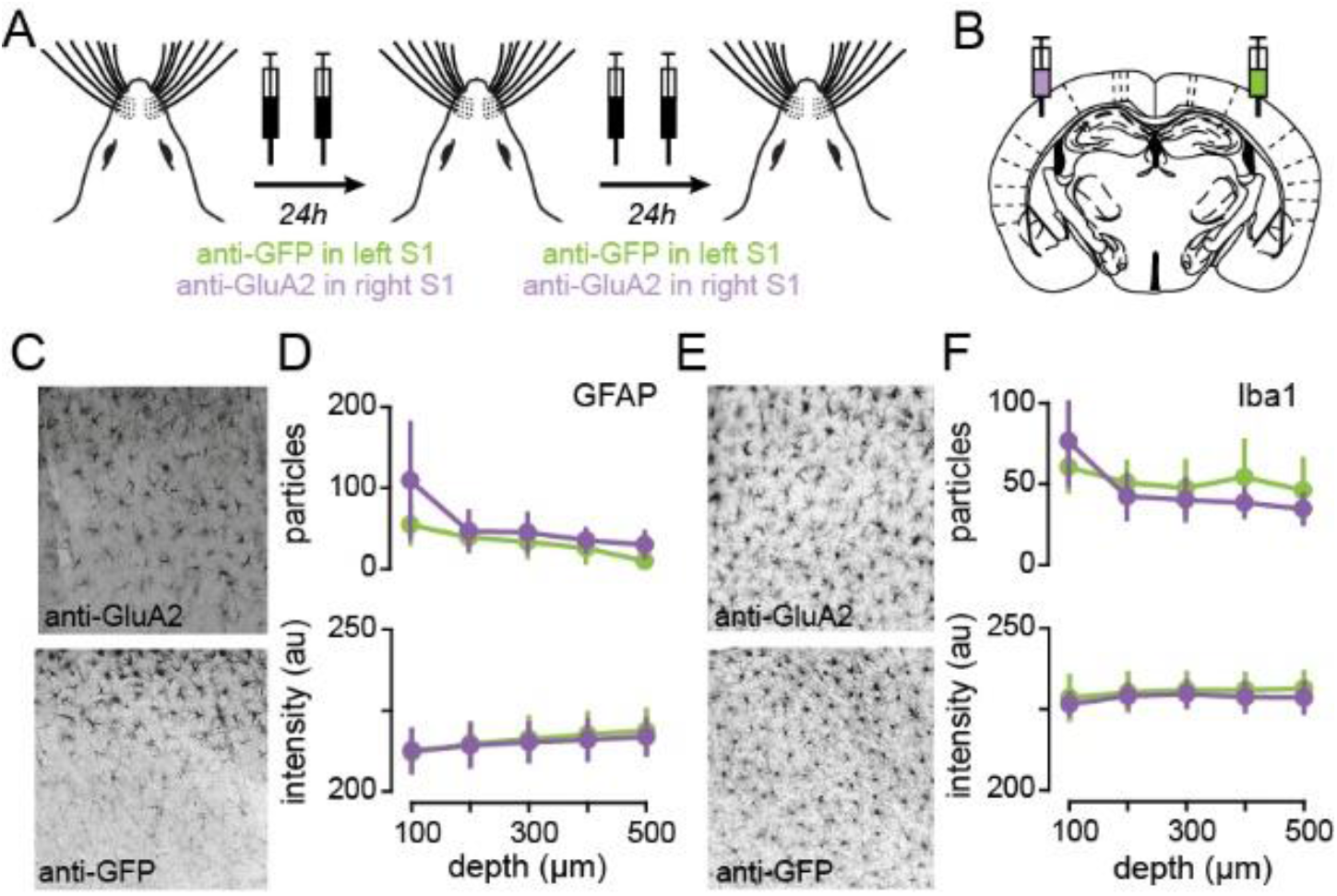
Chronic cross-linking of AMPARs does not increase activated astrocytes nor microglia cells. **A)** Schematic of experimental strategy. IgGs are injected twice a day for two consecutive days in FWE mice. **B)** Anti-GluA2 and anti-GFP IgGs were injected in the right and left hemispheres, respectively. **C)** Immunostaining for the astrocyte marker GFAP, in presence of anti-GluA2 (top) and anti-GFP (*bottom*) IgGs. **D)** Quantification of immunopositive cells (top) and total pixel intensity (bottom) as a function of cortical depth. **E, F)** Same presentation as in (**C, D**) but for the microglia marker Iba1. ***Values and statistical tests are provided in Table S1*.**

**Fig. S6.**
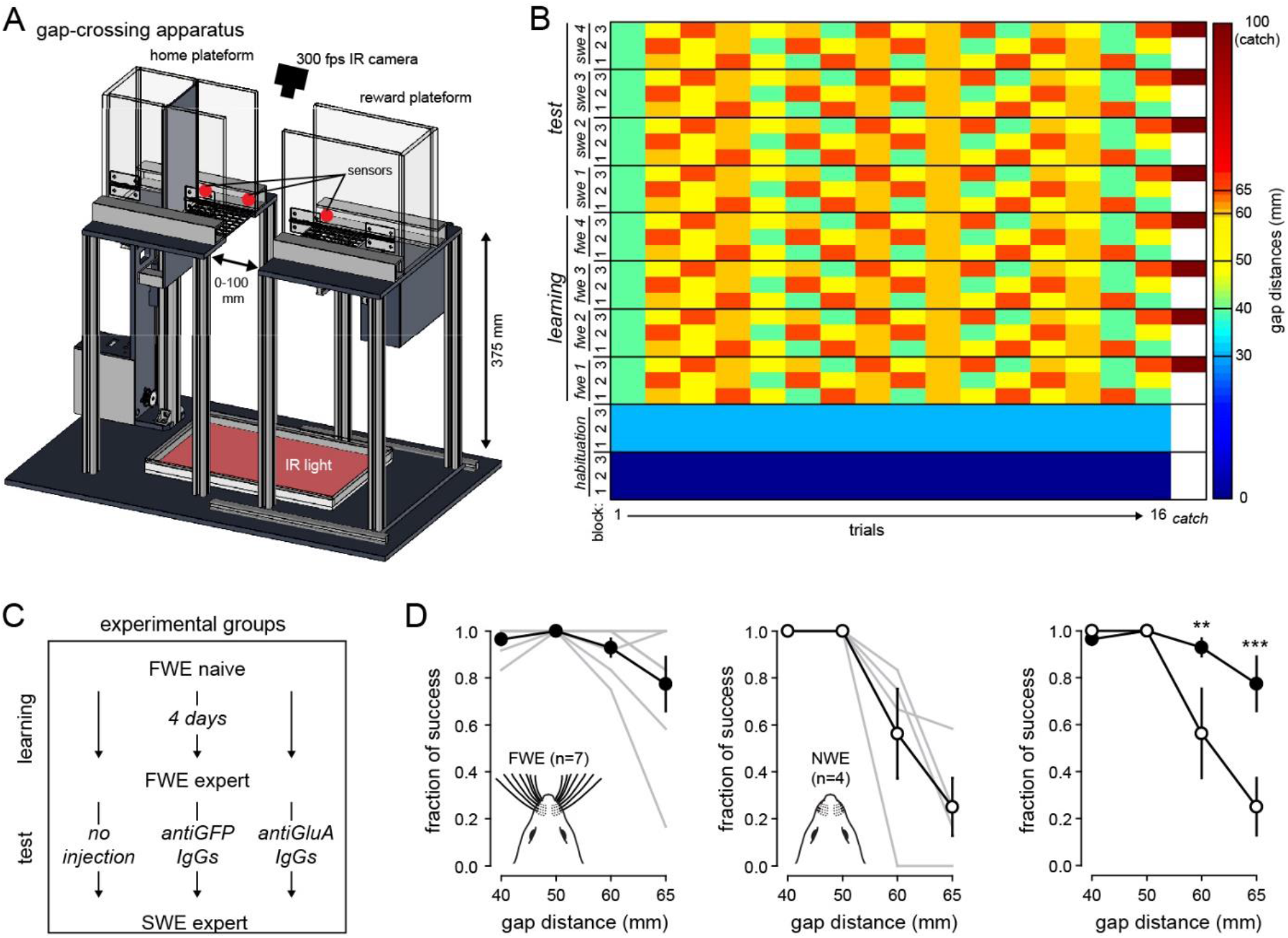
The gap-crossing task relies preferentially on sensory input from whiskers. **A)** Overview of the gap-crossing apparatus. It consists of two individual moveable platforms: (i) a starting platform containing an automated door to precisely control the start of a trial; (ii) a reward platform containing a pellet distributor to deliver a calibrated food reward. Both platforms are elevated 374 mm from the surface and surrounded with 20-cm-high Plexiglas walls. The two platforms face each other with a high-speed 300 fps camera at the top and an infra-red pad at the bottom. The edges of the platforms close to the gap (10 × 10 cm) are made of a metal grid to allow a better grip during jump. A ruler placed in between the platforms is used to precisely define the gap distances (GD) at a given trial. Behavior is done without any sensory cues forcing mice to use their whiskers. **B, C)** Behavioral protocol. Food-restricted mice are first habituated to the apparatus. During test, each session consists of 3 blocks of 16 trials with pseudo-randomized GD (40, 50, 60, and 65 mm). A given trial is defined as success if mice reach the reward platform and eat the food pellet or as a failure if it takes more than 2 min to do so. At the end each trial, the animal is placed back in the home platform to start the next one. Each session ends with a catch trial where the reward platform is removed. This allows to rule out any motor habituation during jumping decision. **D)** Averaged (± sem) fraction of gap-crossing success at a distance of 65 mm, in FWE (*left*) and fully-deprived (no whiskers, NWE, *middle*). Gray lines, individual mice. *Right*, Averaged (± sem) fraction of gap-crossing success at a distance of 65 mm, in FWE (filled circles) and NWE (open circles) mice. ***Values and statistical tests are provided in the Table S1*.**

**Fig. S7.**
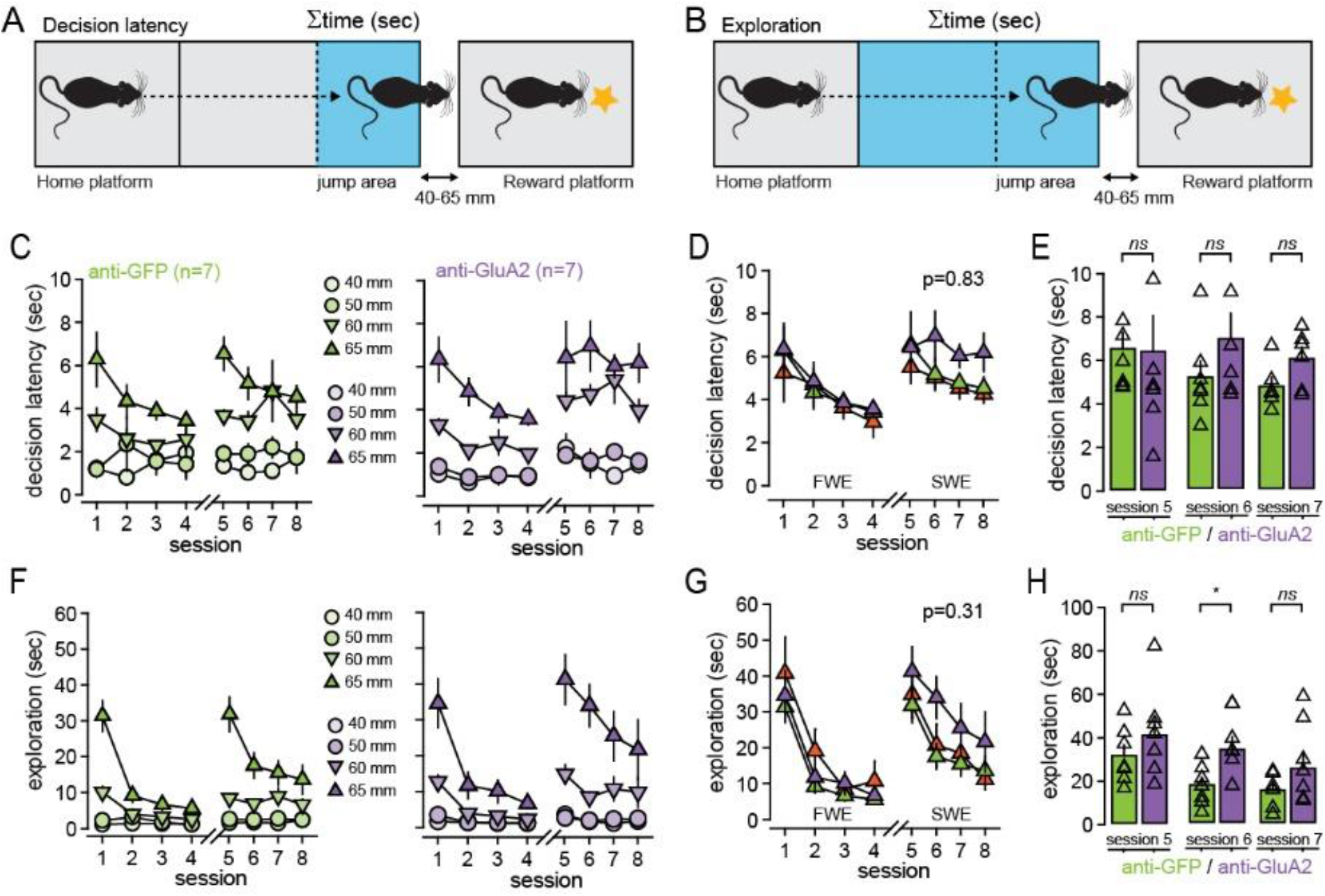
IgGs do not alter exploration and decision latency. **A, B)** Behavioral parameters. The total time (∑time, sec) spent in the jump area (light blue in A) and in the apparatus (light blue in **B**, excluding the start zone) are used as metrics for decision latency and exploration, respectively. **C)** Averaged (± sem) decision latency (sec) for different gap distances, in anti-GFP (*left*) and anti-GLuA2 (*right*) injected mice. **D)** Averaged (± sem) decision latency (sec) at a distance of 65 mm, in non-injected (orange), anti-GFP (green) and anti-GLuA2 (purple) injected mice. **E)** Mean (± sem) decision latency (sec) at a distance of 65 mm after expertise in FWE mice and during SWE. Triangles, individual mice. **F-H)**, Same representation as in **c-e** but for exploration. ***Values and statistical tests are provided in the Table S1.***

**Table S1.**
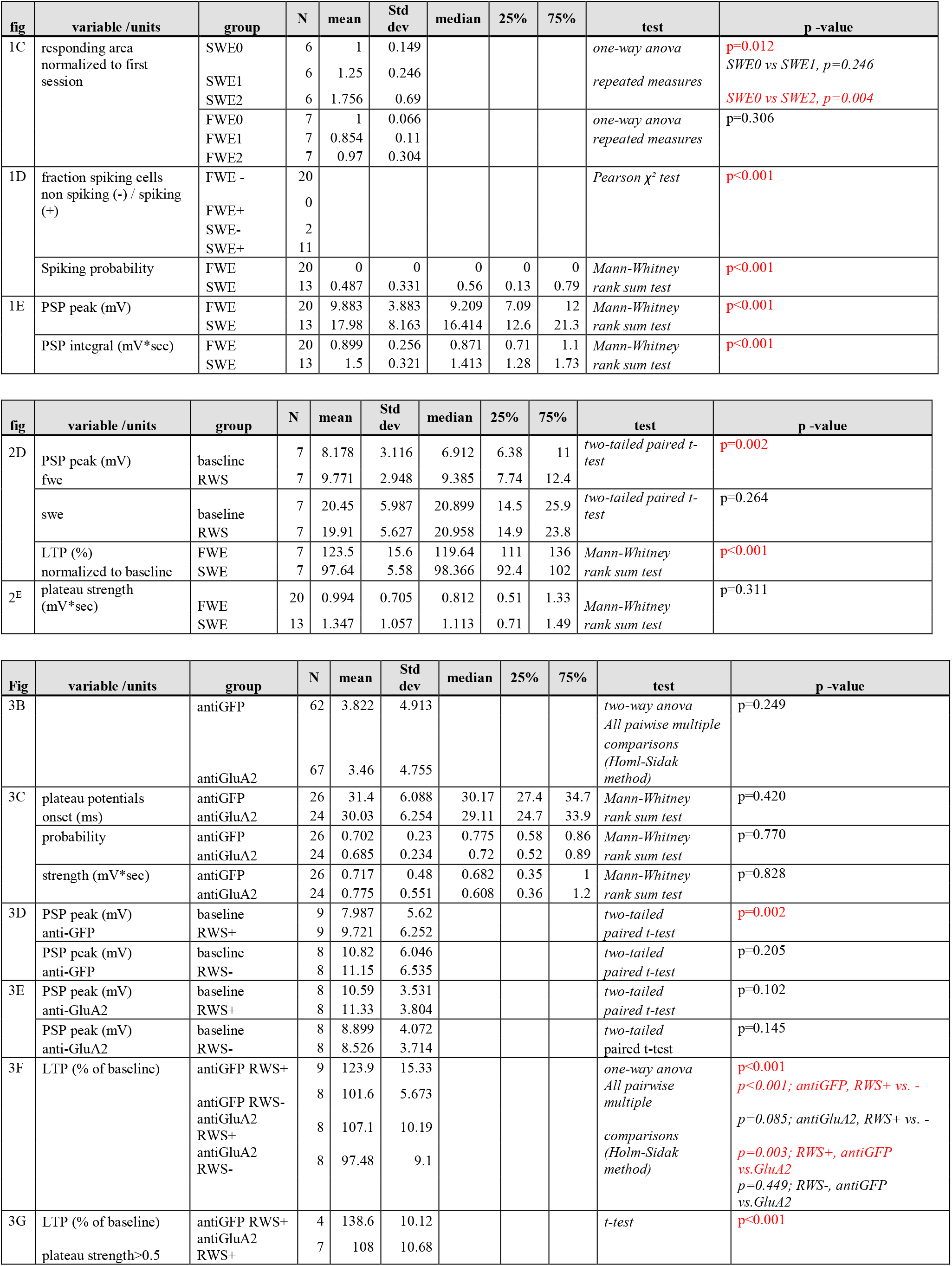
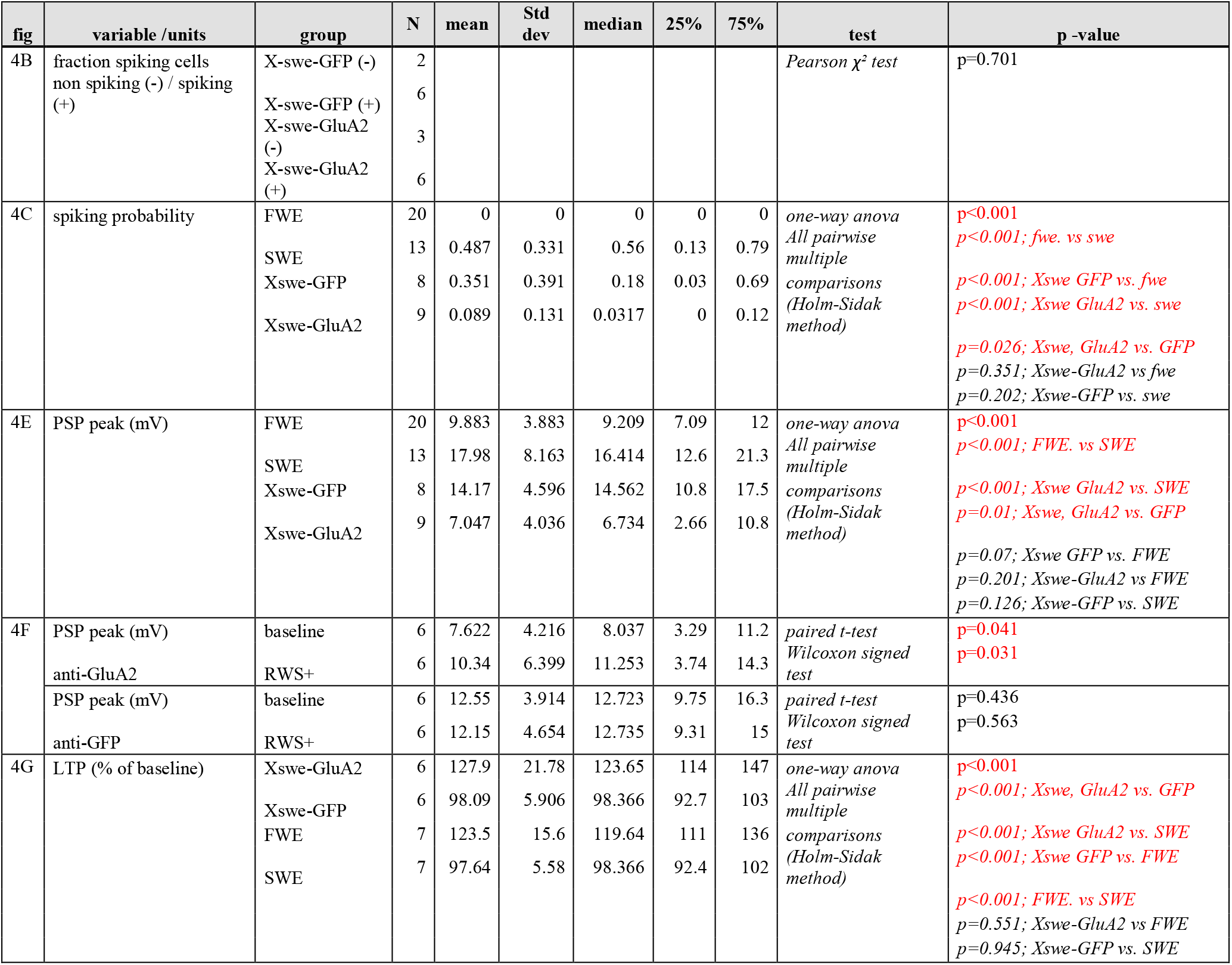
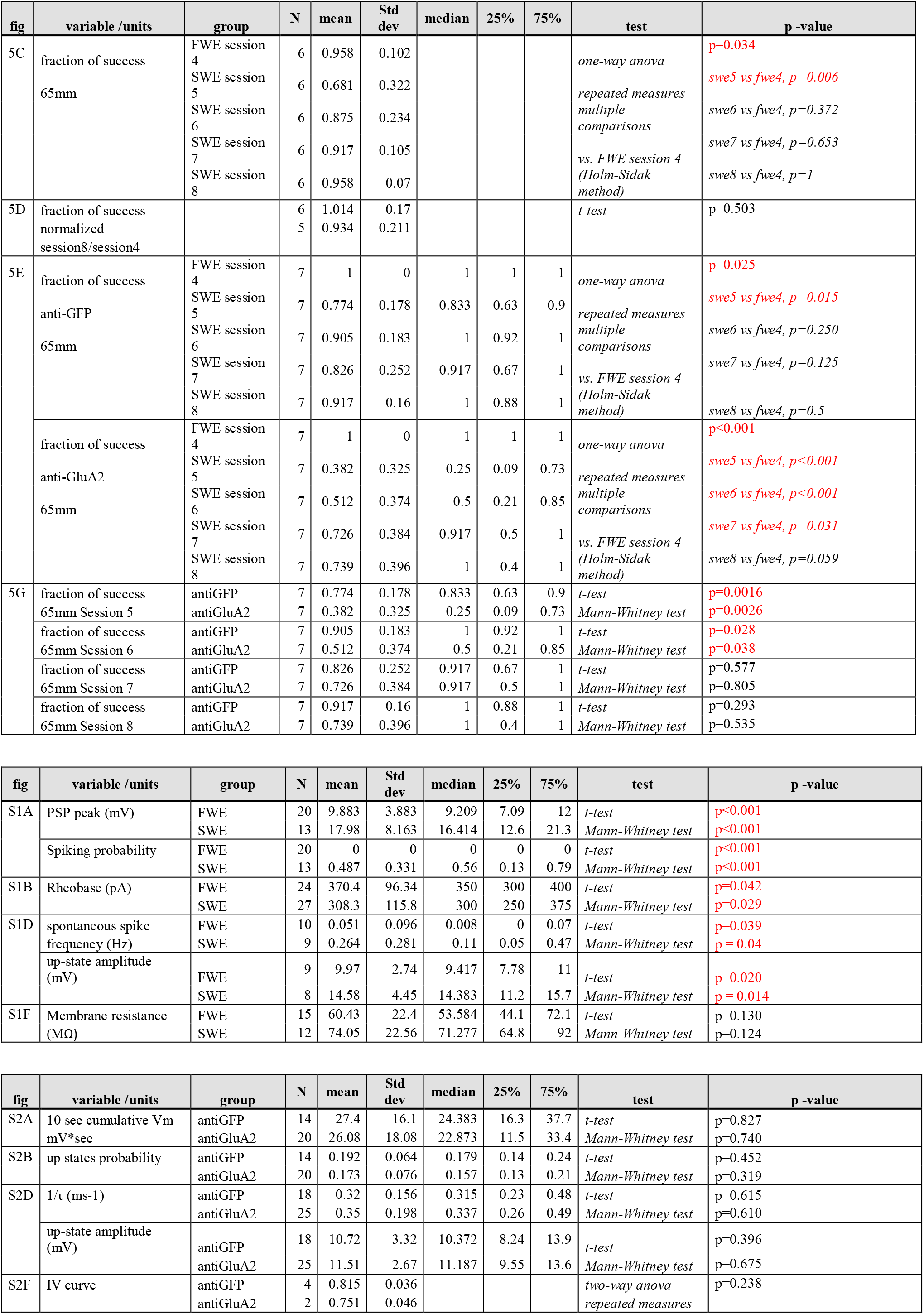
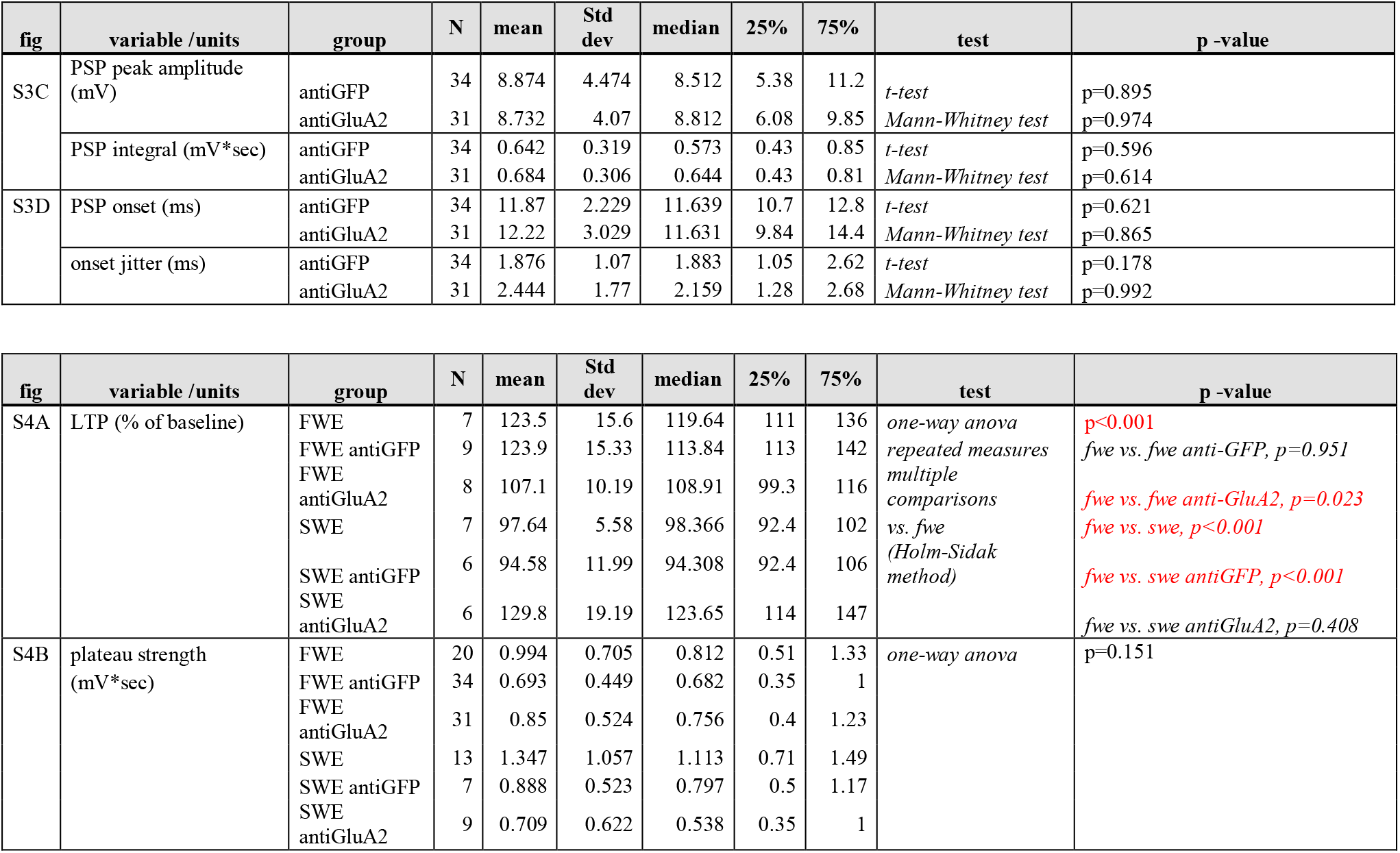
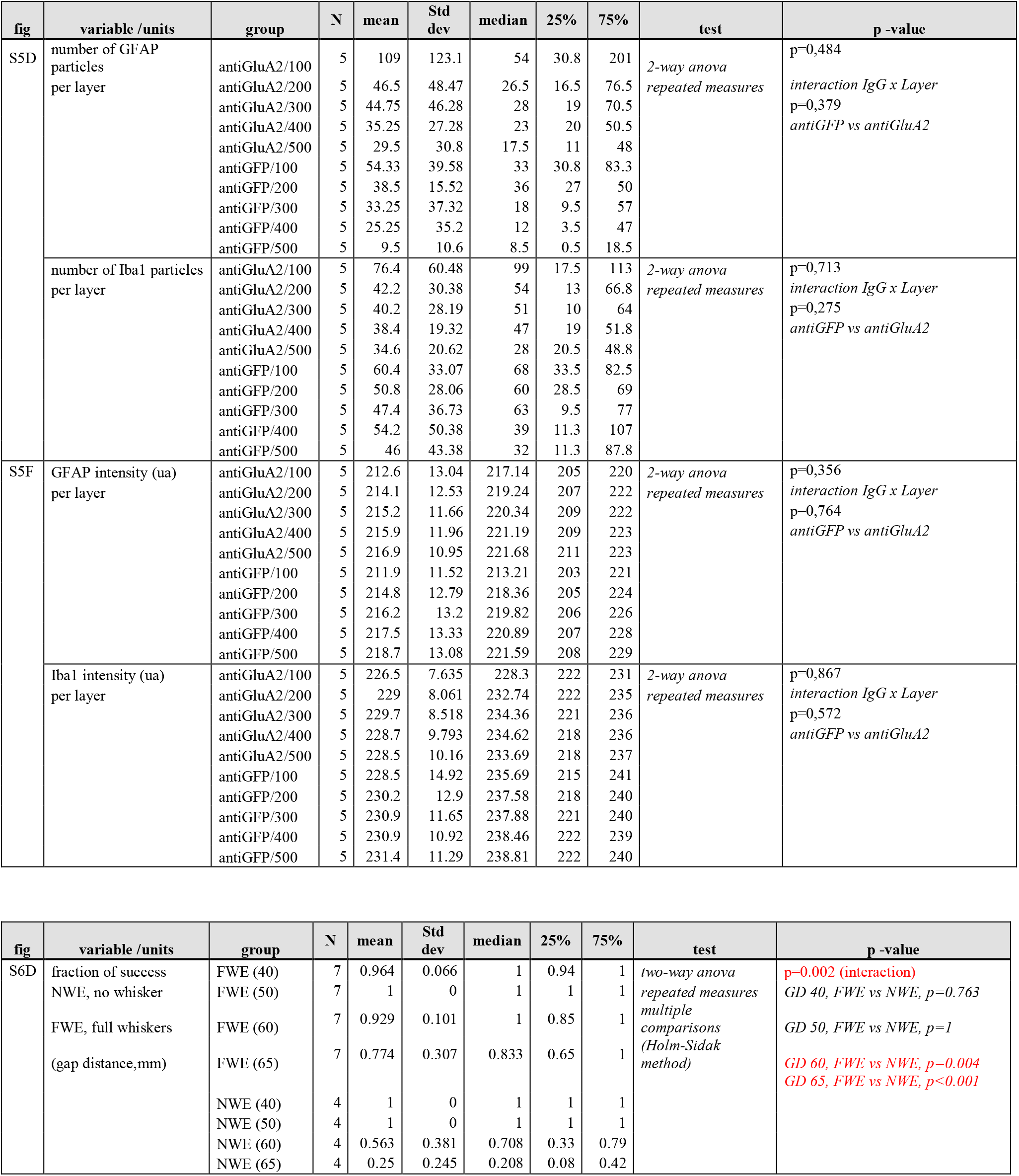
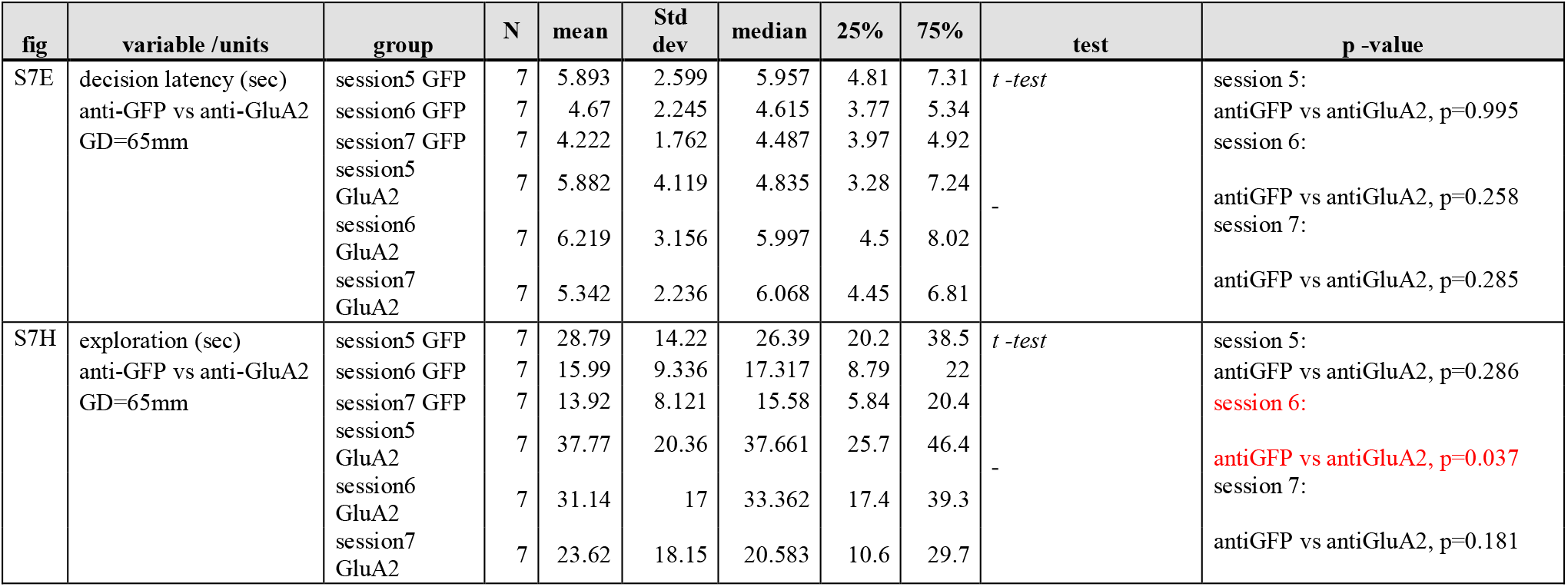

## Methods

### Animals

All experiments were performed in accordance with the Guide for the Care and Use of Laboratory Animals (National Research Council Committee (2011): Guide for the Care and Use of Laboratory Animals, 8th ed. Washington, DC: The National Academic Press.) and the European Communities Council Directive of September 22th 2010 (2010/63/EU, 74). Experimental protocols were approved by the institutional ethical committee guidelines for animal research (N°50DIR_15-A) and by the French Ministry of Research (agreement N°18892). We used male C57BL6/J 5- and 6-weeks old mice from Charles River that were housed with littermates (3 mice per cage) in a 12-h light-dark cycle. Cages were enriched with tunnels, food and water were provided *ad libitum*, except during behavioral experiments (see below).

### Cranial window implantation for chronic Intrinsic Optical Imaging

#### Anesthesia

Anesthesia was induced using isoflurane (4% containing ~0.5 l/min O2) and then continued using an intraperitoneal (i.p.) injection of a MB mixture (MB) (5 μl/g) composed of medetomidine (0.2 mg/kg), and buprenorphine (0.2 mg/kg). A heating-pad was positioned underneath the animal to keep the body temperature at 37°C. Eye dehydration was prevented by topical application of ophthalmic gel. Analgesia was achieved by local application of 100 μL of lidocaine (lurocaine, 1 %) and subcutaneous (s.c.) injection of buprenorphine (buprécare, 0.05 mg/kg). To prevent risks of inflammation and brain swelling 40 μL of dexamethasone (dexadreson, 0.1 mg/mL) were injected intramuscularly (i.m.) before the surgery. After disinfection of the skin (with modified ethanol 70% and betadine), the skull was exposed and a ~5mm plastic chamber was attached to it above the relative stereotaxic location of the C2 barrel column (−1.5 mm from bregma, + 3.3 mm mideline) using a combination of super glue (Loctite) and dental cement (Jet Repair Acrylic, Lang Dental Manufacturing). The chamber was filled with saline (0.9% NaCl) and sealed with a glass coverslip.

#### Intrinsic Optical Imaging (IOI) for barrel column targeting

To locate the cortical barrel column computing the whisker C2 (wC2), intrinsic optical signals (IOS) were imaged as previously described, through the intact skull using a light guide system with a 700 nm (bandwidth of 20 nm) interference filter and stable 100-W halogen light source (*1*–*3*). Briefly, the head of the animal was stabilized using a small stereotaxic frame and the body temperature kept constant with a heating pad. An image of the surface vascular pattern was taken using a green light (546 nm-interference filter) at the end of each imaging session. Images were acquired using the Imager 3001F (Optical Imaging, Mountainside, NJ) equipped with a large spatial 602 × 804 array, fast readout, and low read noise charge-coupled device (CCD) camera. The size of the imaged area was adjusted by using a combination of two lenses with different focal distances (upper lens: Nikon 135 mm, f2.0; bottom lens: Nikon 50 mm, f1.2). The CCD camera was focused on a plane 300 μm below the skull surface. Images were recorded at 10 Hz for 5 sec., with a spatial resolution of 4.65 μm/pixel comprising a total area of 2.9 × 3.7 mm^2^. wC2 was deflected back and forth (20 stimulations at 8 Hz for 1 sec.) using a glass-capillary attached to a piezoelectric actuator (PL-140.11 bender controlled by an E-650 driver; Physik Instrumente) triggered by a pulse stimulator (Master-8, A.M.P.I.). Each trial consisted of a 1 sec. of baseline period (frames 1-10), followed by a response period (frames 11-22) and a post-stimulus period (frames 23-50). Inter-trial intervals lasted 20 sec. to avoid contamination of the current IOS by prior stimulations. IOS were computed by subtracting each individual frame of the response period by the average baseline signal. The obtained IOS was overlapped with the vasculature image using ImageJ software to precisely identify the cortical region computing wC2.

#### Craniotomy and cranial window implantation

After IOI, adequate anesthesia was assessed (absence of toe pinch reflexes, corneal reflexes, and vibrissae movement) and prolonged using supplementary isoflurane if necessary. Dehydration was prevented by injecting sterile saline by s.c. injection. A 3 mm diameter craniotomy was then made over the maximum IOS using a pneumatic dental drill. The craniotomy was covered with sterile saline and sealed with a 3 mm glass coverslip. The coverslip was sealed to the skull using dental acrylic and dental cement (Jet Repair Acrylic, Lang Dental Manufacturing). Anesthesia was reverted by a sub-cutaneous injection of an AB mixture (AB) containing atipamezole (Revertor, 2.5 mg/kg), and buprenorphine (Buprécare, 0.1mg/kg). A delay of 2-3 weeks for surgery recovery was respected before all imaging experiments, during which the body weight of mice was daily checked.

### Chronic Intrinsic Optical Imaging

#### Imaging protocol

MB-anaesthetized mice were daily-imaged during 1 session with all their whiskers (baseline), followed by 2 sessions with all their whiskers trimmed except wC2 (SWE 1-2). A cohort group was additionally recorded for 3 days with all their whiskers (FWE 1-3) as a control for barrel expansion. During each session, wC2 was deflected back and forth (20 stimulations at 8Hz for 1 sec) and IOS recorded through a CW.

#### Spatiotemporal analysis of IOS

An average of 200 trials were recorded per session to quantify IOS as previously described (3). The IOS of different sessions from the same animal were spatially aligned using the animal’s brain surface vasculature and spatially binned (6×6, final resolution: 27.9 μm/pixel or 3×3, final resolution: 13.95 μm/pixel). A high pass-filter was then applied by subtracting from each image-frame the same image-frame that was convolved using a 1270 μm full-width at half maximum (FWHM) Gaussian kernel. The whisker-evoked IOS were then simulated using a pixel-by-pixel paired t-test, comparing the baseline period and the response period of all trials within a session. The t maps for each individual trial were low pass-filtered with a 340 μm FWHM Gaussian kernel and averaged into a final t map response. A threshold was set to t < −2.0 and any signal below this value was considered to belong to the stimulus-evoked response area. If the pixel value was t ≥ −2.0 it was considered background noise and discarded for quantification. This usually resulted in an image with a clear minimum, representing the response maximum and the barrel’s center of mass. Changes on IOS pixel area caused by whisker trimming were computed as the ratio between the whisker-evoked IOS response of the baseline and SWE sessions. All data analysis was performed using a custom software written in MATLAB (MathWorks).

### *In vivo* whole-cell recordings

#### Acute AMPAR X-linking surgery

Anesthesia was induced using isoflurane (4% with 0.5 l min^−1^ O2) and then continue using i.p. injection of urethane (1.5 g kg^−1^). Surgery preparation and IOI were performed as aforementioned. After imaging, adequate anesthesia was assessed and prolonged by supplementary urethane (0.15 g kg^−1^) if necessary. A small ~1 × 1 mm craniotomy (centered above the C2 whisker maximum IOS response) was made using a pneumatic dental drill. Thee injections of either an anti-GluA2 antibody (clone 15F1, gift from E. Gouaux) or a monoclonal anti-GFP IgG1-K (Roche, 11814460001) were targeted to the L2/3 of S1 (−0.1 to 0.3 mm dorsoventral). A 30 nL solution containing antibody diluted in sterile saline (0.05 mg/mL) was injected at maximum rate of 15nl/min, with 30 sec intervals between injection sites as described before. All the experiments were performed blind for the antibody injected.

#### Chronic AMPAR X-linking surgery

Anesthesia was induced using isoflurane (4% containing ~0.5 l min^−1^ O2) and continued using an i.p. injection of MB to perform IOI targeting of the wC2 cortical barrel. Adequate anesthesia was assessed and prolonged using isoflurane if necessary. Dehydration was also prevented by s.c. injection of sterile saline. A small ~ 1 mm diameter craniotomy above the maximum IOS was made using a pneumatic dental drill. The dura was left intact and a stereotaxic injection of either anti-GluA2 or anti-GFP antibody was performed as mentioned above for acute injection. After stereotaxic injection, the craniotomy was covered with sterile saline and protected with a 3 mm polydimethylsiloxane (PDMS) cover slip. PDMS was attached to the skull using an ultra-violet (U.V.) curing optical adhesive (NOA61, Norland) cured with a 50 mW U.V. laser (3755B-150-ELL-PP, Oxxius). Before reverting anesthesia using AB, all the whisker except C2 were trimmed (SWE1). Antibodies were re-injected twice on the day after (SWE2), with a 12h interval between injections using isoflurane anesthesia (4% for induction, then 2% for injection with ~0.5 l min−1 O^2^). Stereotaxic injections were performed through the PDMS cover slip with the same injection protocol than before. After 12h of antibody washout (SWE3), mice were finally anesthetized with isoflurane (4% with 0.5 l min-1 O2) and an i.p. injection of urethane (1.5 g.kg^−1^). Before the patch-clamp recordings, the PDMS C.W. was removed and the cortex protected with sterile saline. All the experiments were performed blind for the antibody injected.

#### Recordings

Whole-cell patch-clamp recordings of L2/3 pyramidal neurons were obtained as described previously (*4*). Current-clamp recordings were made using a potassium-based internal solution in mM: 135 potassium gluconate, 4 KCl, 10 HEPES, 10 Na2-phosphocreatine, 4 Mg-ATP and 0.3 Na-GTP), pH adjusted to 7.25 with KOH, 285 mOsM). High positive pressure (200–300 mbar) was applied to the pipette (5–8 MΩ) to prevent tip occlusion. After passing the pia the positive pressure was immediately reduced to prevent cortical damage. The pipette was then advanced in 1-μm steps, and pipette resistance was monitored in the conventional voltage clamp configuration. When the pipette resistance suddenly increased, positive pressure was relieved to obtain a 3–5 GΩ seal. After break-in, membrane potential (Vm) was measured, and dialysis could occur for at least 5 min before deflecting the whisker. Spiking pattern of patched cells was analyzed to identify pyramidal neurons. Action potentials were obtained by a step-increment of injected current. Spontaneous slow-have fluctuations of the resting membrane potentials were recorded as previously described (*5*). PSPs were evoked by back and forth deflection of the whisker (100 ms, 0.133 Hz) as previously described (*4*). The voltage applied to the actuator was set to evoke a displacement of 0.6 mm with a ramp of 7-8 ms of the wC2. Different frequencies of stimulation were used accordingly to the experiment (RWS-LTP: 8Hz, 1 min; cumulative PSPs: 8Hz, 2.5 sec). Series and input resistance were monitored with a 100-ms long-lasting hyperpolarizing square pulse 400 ms before each single-deflection and extracted offline by using a double-exponential fit. Recordings were discarded if the change in these parameters were larger than 30%. The bridge was usually not balanced, and liquid junction potential not corrected. All the data were acquired using a Multiclamp 700B Amplifier (Molecular Devices) and digitized at 10 kHz (National Instruments) using software. Offline analysis was performed using custom routines with IGOR Pro (WaveMetrics).

### Behavior

#### Gap crossing apparatus

The custom-made gap crossing (G.C.) apparatus (Imetronic, France) consists of two individual moveable platforms: (1) a starting platform containing an automated door to precisely control the start of a trial; (2) a reward platform containing a pellet distributor to deliver a calibrated food reward. Both platforms (10×20 cm) were elevated 37.4 cm from the surface and surrounded on the three sides with a 20-cm-high Plexiglas walls. The two platforms were placed facing each other with a high-speed 300 frames per second (fps) camera at the top and an infra-red pad at the bottom. This allowed us to precisely track mice behavior and whisker motion with high spatiotemporal resolution. The edges of the platforms that face each other were made of a metal grid (10 × 10 cm) to allow a better grip where the animals should jump. A ruler placed at the bottom and between the platforms was used to precisely define the gap distances (GD) at a given trial. The apparatus was placed into a ligh- and proof cage containing ventilation, and surrounding speakers with a continuous white noise background. This ensures that mice do not have neither visual nor auditory cues regarding the reward platform. Food pellet odor was saturated inside the box to avoid any olfactory-related cues.

#### Behavioral protocol

At least 5 days before starting behavior, mice were food restricted and handled to decrease stress. After a 15 – 20 % reduction of the initial body weight, habituation was performed during 3 days: (day 1 – Maze Habituation) mice were placed on the G.C. apparatus with a GD = 0 cm for 10 min. where the pellet distributor was randomly presented for multiple times without food reward; (day 2 – Reward Habituation) mice were placed on the start platform and trained for 3 blocks (16 trials each block, GD = 0 cm) to the distribution of a food pellet in the reward platform. A given trial was defined as success if the animal reached the reward platform and ate the food pellet or as a failure if it took more than 2 min to do so. At the end of a trial, the animal was placed back in the starting platform to beginning the next one; (day 3 – Jump Habituation) the same protocol than (2) but using a GD = 3 cm to habituate the animal for a distance between platforms. Habituation is considered successful and the test sessions started if the success rate was >95%. The test protocol had 1 session per day during 4 days where each session was composed of 16 trials containing GD = 40, 50, 60, and 65 mm. Individual blocks started with the minimal GD, had random GD sequences, and finished with a catch trial (GD: 100 mm) where the reward platform was removed. This allowed to rule out habit to jump. When addressing the effect of whisker trimming on expert mice, test sessions were performed before and after whisker trimming.

#### Cannula implantation for chronic AMPAR X-linking

Anesthesia was induced using isoflurane and continued by an i.p. injection of MB to perform IOI targeting of the wC2 cortical barrel as aforementioned. A small craniotomy above the maximum IOS was made using a pneumatic dental drill, preventing any cortical damage. After drilling, a guide cannula (62001, RWD Life Science Co., LTD) was stereotaxically inserted in the brain using a cannula holder through the craniotomy previously made. The size of the cannula (0.6 mm) was adjusted to target L1 of the somatosensory cortex. The guide cannula was fixed to the skull using two stainless steel screws and a mix of super glue (Loctite), dental acrylic and dental cement. Anesthesia was reverted by a s.c. injection of AB and mice left to recover over 2 weeks before starting food restriction. During food restriction, mice were additionally habituated to be restrained by a different experimenter to avoid stress during antibody injection. Mice were tested during 4 sessions with FWE followed by 4 SWE sessions, during which either an anti-GluA2 or an anti-GFP antibody (0.05 mg/mL) was injected. Antibodies were injected twice per day, before and after each test session, using a pump (D404, RWD Life Science CO.) with an injection speed of 6nL/min for the first 120nL and 3nL/min for the remaining 30nL of antibody. Mice were freely moving in their home cage during injection. All experiments and analysis were performed blind for the antibody injected.

### Histology

To evaluate the antibody injection profiles in S1, animals were intracardially perfused with PBS (1%) and PFA (4%). Fixed brains were sliced with a vibratome and sections incubated with PBS.H202 (0.3%) during 30 min to block endogenous peroxide. Brain slices were then incubated with a secondary anti-mouse biotinylated antibody from donkey (1/200), during 2h at room temperature (RT). To finally reveal the injected primary antibody, slices were first incubated with an avidin-biotin complex (1/200 in PBS (1×) – Triton 0.1%), and then with DAB (ab64259, Abcam). Brain slices were finally mounted between slide and coverslip and imaged post-hoc using a Nanozoomer (S360, Hamamatsu). Illumination was set such that the full dynamic range of the 16-bit images was utilized. A two-dimensional graph of the intensities of pixel was then plotted using a Fiji Software. 16-bit image’s brightness was processed and a mask was registered to the corresponding coronal plates (ranging from −0.26 to −1.94 mm) of the mouse brain atlas using Illustrator (Adobe), at the various distances posterior to the bregma.

To evaluate the astrocytic and microglial reactivity, anesthesia was induced using isoflurane (4% containing ~0.5 l min^−1^ O2) and continued by adjusting its percentage to 1.5-3%. One small ~1 × 1 mm craniotomy was performed in each hemisphere, targeting the barrel field (Fronto-caudal: −1.0 / Mediolateral: ± 3.3). The dura was left intact to then perform stereotaxic injection of either an anti-GluA2 (left hemisphere) or anti-GFP antibody (right hemisphere). After stereotaxic injection the craniotomy was covered with sterile saline and protected with a 3 mm polydimethylsiloxane (PDMS) cover slip as previously described. Antibodies were re-injected twice the day after surgery, with a 12h interval between injections using isoflurane anesthesia (4% for induction, then 2% for injection with ~0.5 l min^−1^ O2). After 12h of antibody washout, mice were finally anesthetized with isoflurane (4% with 0.5 l min^−1^ O2) and a mix of pentobarbital sodium and lidocaine (300 mg/kg; 30 mg/kg; i.p.) for animal’s intracardiac perfusion with PBS (1%) and PFA (4%). Fixed brains were sliced with a vibratome and sections were incubated afterwards in PBS with Triton X-100 0.3% for 2h. A subset of brain slices were incubated either with an anti-GFAP antibody (chicken anti-GFAP, G2032-25F-100ul, USBiological Life Sciences) or an anti-Iba1 (rabbit anti-Iba1, 019-19741, WAKO) antibody (1/500 in PBS-Tween 0.05%) overnight at 4C°. Primary antibodies were washed-out 3 times with PBS, and secondary antibodies (Alexa 647 anti-chicken: A32933, Alexa 647 anti-rabbit: A27040, Thermofisher, 1/500 in PBS-Tween 0.05%) incubated at RT for 3h. Brain slices were finally mounted between slide and cover slip and imaged post-hoc using a Nanozoomer (S360, Hamamatsu). Importantly, the same settings of imaging were kept for all the recorded samples.

Raw data were converted to 8-bit images on ImageJ and segmented with ROI’s with similar width and height corresponding to 100 μm to determine the staining across all the superficial cortical layers. An average of pixel intensity for each ROI was calculated to determine staining intensity. LUT’s were then inverted and fitted with a Gaussian blur (5 pixels) to smooth the image and reduce noise. Finally, an entropy threshold (https://imagej.net/Maximum_Entropy_Threshold, ImageJ) was applied and the resulting number of particles for each individual ROI calculated. This allowed us to calculate the number of GFAP- or Iba1-positive cells across cortical layers.

### Statistics

Detailed statistics are described in Extended Data Table 1. Statistical differences were considered at p<0.05. All experiments and analysis were performed blind for experimental conditions.

